# Modeling microbiome-trait associations with taxonomy-adaptive neural networks

**DOI:** 10.1101/2023.11.04.565596

**Authors:** Yifan Jiang, Matthew Aton, Qiyun Zhu, Yang Young Lu

**Author notes:** Corresponding author(s). E-mail(s); Contributing authors.

## Abstract

The human microbiome, a complex ecosystem of microorganisms inhabiting the body, plays a critical role in human health. Investigating its association with host traits is essential for understanding its impact on various diseases. Although shotgun metagenomic sequencing technologies have produced vast amounts of microbiome data, analyzing such data is highly challenging due to its sparsity, noisiness, and high feature dimensionality. Here we develop MIOSTONE, an accurate and interpretable neural network model for microbiome-disease association that simulates a real taxonomy by encoding the relationships among microbial features. The taxonomy-encoding architecture provides a natural bridge from variations in microbial taxa abundance to variations in traits, encompassing increasingly coarse scales from species to domains. MIOSTONE has the ability to determine whether taxa within the corresponding taxonomic group provide a better explanation in a data-driven manner. MIOSTONE serves as an effective predictive model, as it not only accurately predicts microbiome-trait associations across extensive simulated and real datasets but also offers interpretability for scientific discovery. Both attributes are crucial for facilitating *in silico* investigations into the biological mechanisms underlying such associations among microbial taxa.

## 1 Introduction

The human microbiome characterizes the complex communities of microorganisms living in and on our bodies, with bacteria alone encoding 100 times more unique genes than humans (Qin et al, 2010). As the microbiome influences the impact of host genes, microbiome genomes are often referred to as the “second genome” (Grice and Segre, 2012). Subsequently, microbiomes have been found to play pivotal roles in various aspects of human health and diseases (Sekirov et al, 2010), including diabetes (Qin et al, 2012), obesity (Turnbaugh et al, 2006), inflammatory bowel disease (Mills et al, 2022), Alzheimer’s disease (Vogt et al, 2017), and more. The association of the microbiome with host traits will provide insights into the underlying mechanisms governing the microbiome’s impact on human health and diseases and facilitate the development of novel therapeutic strategies.

To investigate microbiome-trait associations, one primary focus has been on identifying predictive microbial markers for disease prediction from microbial samples (Medina et al, 2022). Here, a microbial sample is typically characterized by its taxonomic profile, which includes the abundance of microbial taxa at certain taxonomic levels (Pasolli et al, 2016), such as species, genus, family, and so on. However, the unique characteristics of microbiome data pose challenges in thoroughly exploring the relationships among the taxa. For example, sample-wise sequencing generates millions of short fragments from a mixture of taxa rather than an individual taxon. The dynamic and complex nature of microbial communities can lead to inaccurate taxonomic profiling, potentially resulting in inaccuracies in downstream microbiome-trait association analyses (Ye et al, 2019).

Another difficulty in analyzing microbiome data stems from its sparsity, with a substantial portion of data entries being zeros (Medina et al, 2022). These zeros can indicate either the true absence of the taxa in the environmental sample (*i.e.,* biological zeros) or the failure to detect the taxa due to low sequencing depth and sampling variation (*i.e.,* technical zeros) (Jiang et al, 2021). To address the sparsity issue, existing imputation methods are typically employed to distinguish technical zeros from biological zeros and replace technical zeros with nonzero values (Jiang et al, 2021; Zeng et al, 2022; Linderman et al, 2022). However, these methods require subjective user decisions to choose the threshold that decide which zeros require imputation and which do not (Zeng et al, 2022), inevitably diminishing the reliability and reproducibility of the downstream analyses. Moreover, imputation methods can lead to data misinterpretation by introducing the risk of bias and potentially yielding false signals (Jiang et al, 2022; Andrews and Hemberg, 2018).

Additionally, microbiome data is typically high-dimensional and noisy, with a much larger number of taxa than sample size (Kharchenko, 2021). The excessive number of taxa as features not only increases computational costs but also presents challenges in analysis due to the curse of dimensionality. (Liu et al, 2017). Specifically, the relatively low number of samples lead to overfitting during training, thereby limiting generalization to other datasets. For example, the small sample size might lead to conflicting results when inferring the association between microbiome and disease states (Knights et al, 2011; Finucane et al, 2014). To alleviate the curse of dimensionality, feature selection methods have been developed to choose highly variable taxa (Stuart et al, 2019; Hao et al, 2021; Ditzler et al, 2015). However, these methods often result in a significant loss of information from non-selected taxa (Kharchenko, 2021). Altogether, the nature of microbiome data necessitates novel analytic methods that consider factors such as data imperfections, sparsity, and the curse of dimensionality.

To address these challenges, recent studies have leveraged the inherent correlation structure among taxa as an informative prior, with the aim of enhancing disease prediction performance. Two fundamental correlation structures among taxa are taxonomy and phylogeny (Washburne et al, 2018). The taxonomy categorizes taxa into hierarchical groups, spanning from the three domains (Bacteria, Archaea, and Eukarya) down to species (Parks et al, 2018), using a well-established and widely accepted naming system. The phylogeny aims to encode the evolutionary relationships among taxa and classifies taxa by a series of splits, corresponding to estimated events in which two lineages split from a common ancestor to form distinct species (Washburne et al, 2018). The distinction between taxonomy and phylogeny lies in the fact that taxonomy is coarser, with taxonomic labels categorizing only a small fraction of the branches in the phylogeny, whereas the phylogeny provides a more detailed scaffold. Both taxonomy and phylogeny have been utilized to incorporate relevant structural knowledge among taxa into existing predictive models. For example, phylogeny can serve as a smoothness regularizer to enhance linear regression models (Xiao et al, 2018). Utilizing phylogeny can also aid in weighting and prioritizing the most relevant taxa, thus enhancing the prediction accuracy of random forest models (Albanese et al, 2015). More recently, several methods have incorporated phylogeny or taxonomy into the preprocessing stage of employing advanced analysis tools like deep neural networks (DNN) (Sharma et al, 2020; Reiman et al, 2020; Li et al, 2021; Shtossel et al, 2023; Wang et al, 2021). Specifically, these preprocessing steps involve aggregating taxa into distinct taxonomic clusters (Sharma et al, 2020; Wang et al, 2021), reordering the taxa spatially based on their phylogenetic structure (Reiman et al, 2020; Shtossel et al, 2023), and assigning varying weights according to phylogenetic distances (Li et al, 2021). However, these methods may underutilize the relationships inherent in phylogeny or taxonomy by confining their power to the preprocessing step while relegating the DNN to a black box model. While black box modeling remains useful, it proves insufficient for reasoning about the mechanisms governing dynamic interactions among taxa (Ma et al, 2018) – an aspect that is critical for a scientific understanding of microbiome-trait associations.

In this study, we introduce MIOSTONE (<u>MI</u>cr<u>O</u>biome-trait a<u>S</u>sociations with <u>T</u>ax<u>ON</u>omy-adaptiv<u>E</u> neural networks), an accurate and interpretable neural network model that simulates a real taxonomy by encoding the relationships among microbial features (Fig. 1**a**). Drawing inspiration from biologically-informed DNNs (Ma et al, 2018; Elmarakeby et al, 2021), the model organizes the neural network into layers to explicitly emulate the taxonomic hierarchy within its architecture, based on the Genome Taxonomy Database (GTDB) (Parks et al, 2018), spanning 124 phyla, 320 classes, 914 orders, 2,057 families, 6,811 genera, and 12,258 species. In this taxonomy-encoding network, each neuron represents a specific taxonomic group, with connections between neurons symbolizing the hierarchical subordination relationships among these groups. These hierarchies provide a natural bridge from variations in microbial taxa abundance to variations in traits, encompassing increasingly coarse scales from species to domains. The key novelty of MIOSTONE lies in the unique capability of its internal neurons to determine whether taxa within the corresponding taxonomic group offer a more effective explanation of the trait when considered holistically as a group or individually as distinct taxa. Such taxonomy-adaptive strategy is achieved during the training phase, where variations in microbial taxa propagate individually through the hierarchy to impact the parent taxonomic group that contain them, competing against the aggregation of the parent as a whole for better trait prediction (Fig. 1**b**). The taxonomy-encoding design significantly reduces the model’s complexity, thereby mitigating the curse of dimensionality and overfitting, while also providing a natural interpretation of the model’s internal mechanisms. We have applied MIOSTONE to various tasks across extensive simulated and real datasets to demonstrate its empirical utility (Fig. 1**c**). From a practitioner’s perspective, MIOSTONE serves as an effective predictive model, as it not only accurately predict microbiome-trait associations but also be interpretable. Both attributes are crucial for facilitating *in silico* investigations into the biological mechanisms underlying such associations among

**Fig. 1.**
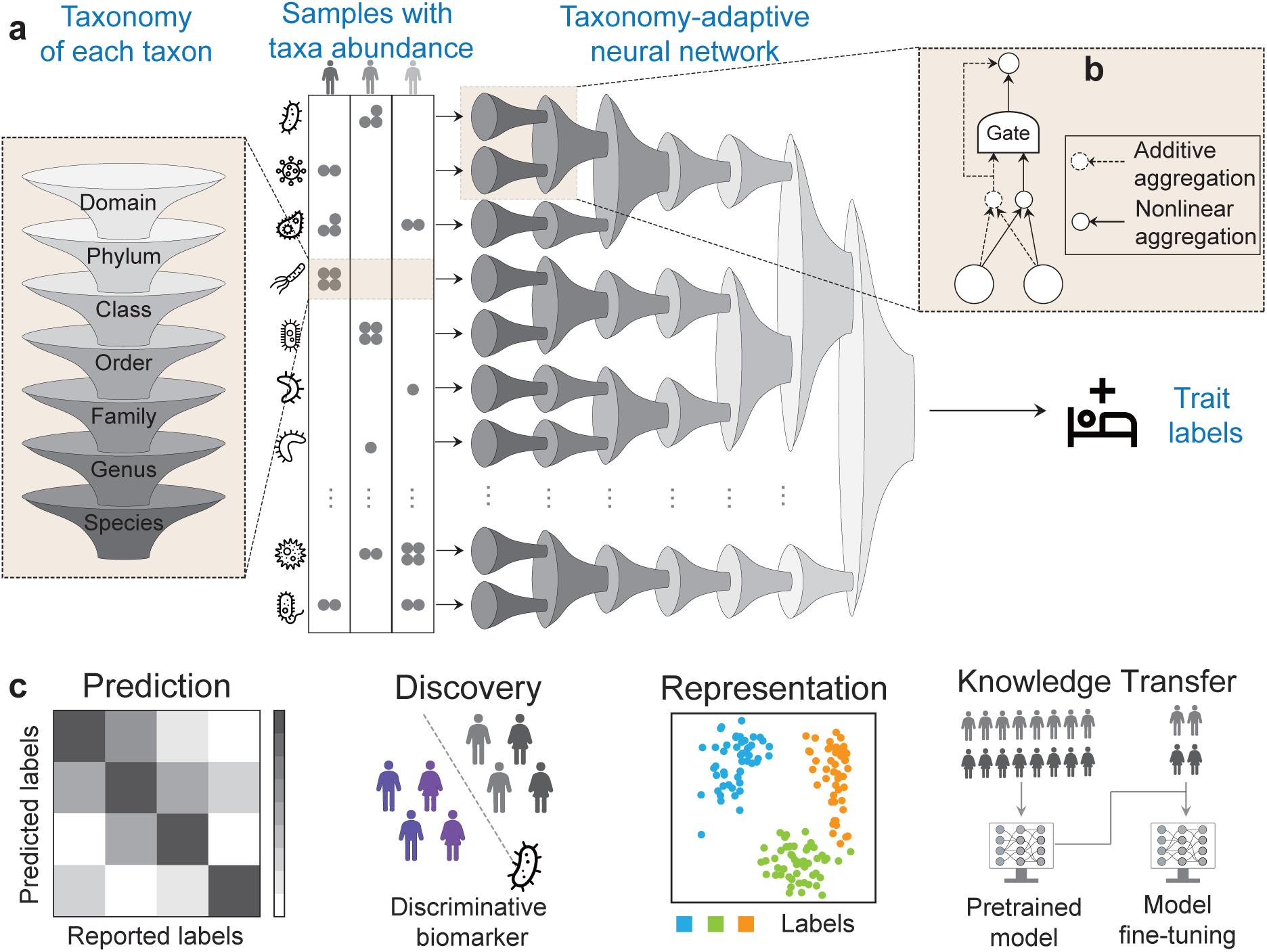
Overview of MIOSTONE. (**a**) MIOSTONE designs the neural network with layered architecture, explicitly mirroring the taxonomic hierarchy. Each neuron represents a distinct taxonomic group, while connections between neurons signify hierarchical subordination relationships among these groups. By contrast, a conventional neural network functions as a black box, lacking explicit encoding of knowledge within its architecture. (**b**) Each internal neuron in MIOSTONE has the capability to discern whether taxa within the corresponding taxonomic group provide a more effective explanation of the trait when assessed either holistically (*i.e.,* additively) as a group or individually (*i.e.,* non-linearly) as distinct taxa. (**c**) MIOSTONE establishes a versatile microbiome data analysis pipeline, applicable to a variety of tasks including disease status prediction, microbiome representation learning, microbiome-disease association identification, and enhancement of predictive performance in tasks with limited samples through knowledge transfer.

microbial taxa.

## 2 Results

### 2.1 MIOSTONE provides accurate predictions of the host’s disease status

We evaluated MIOSTONE’s performance in disease status prediction using three simulated and seven real microbiome datasets. These datasets vary in microbial taxa and sample sizes, encompassing distinctly different taxonomic structures (Fig. A.1). These datasets span various disease prediction tasks, including Autism Spectrum Disorder, Alzheimer’s disease, Graves’ disease, Parkinson’s disease, and inflammatory bowel disease (Refer to Sec. 4.1 and Sec. A.1 for details). Together, they offer a comprehensive evaluation framework for predictive models of the human gut microbiome.

We benchmarked MIOSTONE with the other nine baseline methods, divided into two categories: tree-agnostic methods and tree-aware methods. Tree-agnostic methods, such as Random Forest (RF), Support Vector Machine (SVM), XGBoost, and multi-layer perceptron (MLP), are widely used for predicting disease status (Medina et al, 2022). Notably, tree-aware methods, such as DeepBiome (Zhai et al, 2024), Ph-CNN (Fioravanti et al, 2018), PopPhy-CNN (Reiman et al, 2020), TaxoNN (Sharma et al, 2020), and MDeep (Wang et al, 2021), are specifically designed to leverage phylogenetic or taxonomic structures in microbial taxa to enhance disease prediction. (Refer to Sec. 4.3 and Sec. A.2 for more baseline details).

We assessed the predictive performance of all methods using 5-fold cross-validation, training separate models on each training split and evaluating them on the corresponding test splits. Predictions from all test splits were concatenated, and evaluation was performed on the combined dataset. To quantify predictive performance, we utilized two metrics: the Area Under the Receiver Operating Characteristic Curve (AUROC) and the Area Under the Precision-Recall Curve (AUPRC). For robustness, we repeated this process 20 times with different random seeds and reported the mean performance with 95% confidence intervals. Each repetition involved both data splitting and ML model training.

Our analysis shows that MIOSTONE notably outperformed all tree-agnostic methods and tree-aware methods across all datasets, in terms of AUPRC (Fig. 2) and AUROC scores (Fig. A.2). XGBoost, the second-best baseline method, outperforms MIOSTONE in only three out of ten datasets, with a 3.5% improvement in AUPRC score. However, it performs worse than MIOSTONE in six out of ten datasets, with a 13.7% decrease in performance. Within the tree-aware method category, MIOSTONE outperforms MDeep, the second-best tree-aware method, in eight out of ten datasets with a 3.8% improvement in AUPRC score, while the performance is tied in the remaining two datasets. For scientific rigor, we quantified the performance comparison between MIOSTONE and each baseline method using one-tailed two-sample t-tests to calculate p-values. These p-values confirm that MIOSTONE’s performance superiority is both statistically significant. The superior performance across diverse microbiome datasets and disease models demonstrates MIOSTONE’s robustness and generalizability.

**Fig. 2.**
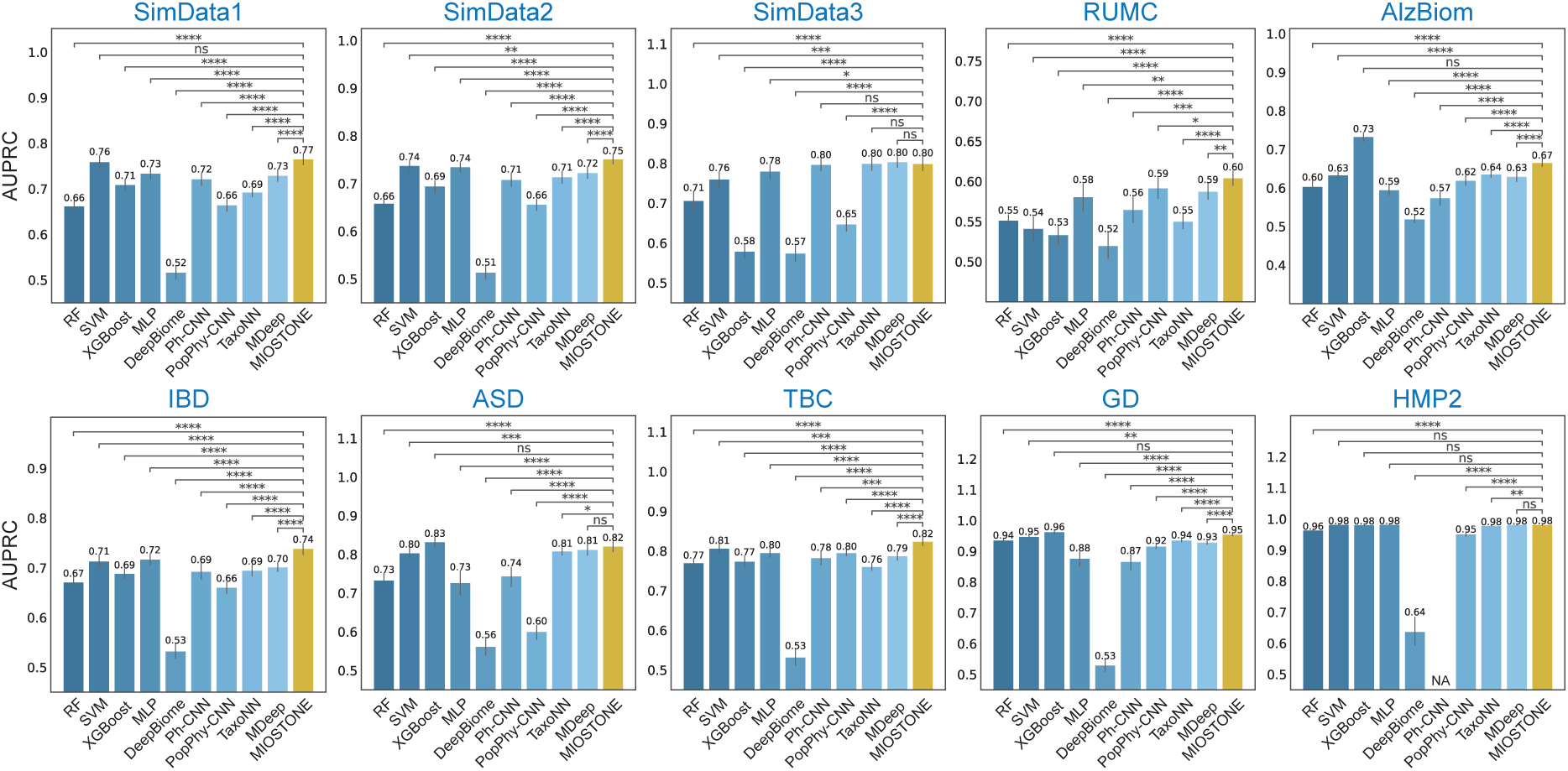
MIOSTONE provides accurate predictions the host’s disease status. The evaluation was performed on three simulated and seven real microbiome datasets with varying microbial taxa sizes, covering different proportions of taxonomic levels. MIOSTONE is compared against nine baseline methods, divided into two categories: tree-agnostic methods and tree-aware methods. The former category comprises random forest (RF), support vector machine (SVM) with a linear kernel, XGBoost, and multi-layer perceptron (MLP), while the latter includes DeepBiome, Ph-CNN, PopPhy-CNN, TaxoNN, and MDeep. Each model was trained by times using different train-test splits, and reported by the average performance along with 95% confidence intervals. The models’ performances are measured by the Area Under the Precision-Recall Curve (AUPRC). Because Ph-CNN is not scalable for processing the HMP2 dataset, the result is denoted as N/A. For scientific rigor, the performance comparison between MIOSTONE and any other baseline method is quantified using one-tailed two-sample t-tests to calculate p-values: ∗ ∗ ∗ ∗ p-value ≤ 0.0001; ∗ ∗ ∗p-value ≤ 0.001; ∗ ∗ p-value ≤ 0.01;∗ : p-value ≤ 0.05; ns : p-value *>* 0.05.

Finally, we observed that model complexity, a key characteristic of tree-aware methods, significantly impacts predictive performance. For example, in datasets like RUMC, higher model complexity improves predictive performance in tree-aware methods compared to tree-agnostic methods. However, in other datasets, such as GD, this increased complexity negatively affects performance, resulting in lower accuracy compared to tree-agnostic methods. Notably, in all examined scenarios, MIOSTONE consistently provides the most accurate predictions of the host’s disease status, striking an optimal balance between model complexity and predictive power.

### 2.2 Dissecting the performance of MIOSTONE

We first evaluated the computational cost of MIOSTONE relative to the other baseline methods (Fig. 3**a**). For a fair comparison, all models were trained and tested in the same environment (Refer to Sec. A.2 for hardware details). Furthermore, all tree-aware models and MLP are trained for a fixed 200 epochs with a consistent batch size of 512 to ensure convergence. It is important to note that the comparison is not entirely fair, as tree-agnostic methods except MLP have their own criteria for determining when to stop training. The results show that MIOSTONE is relatively efficient to train, comparable to training an MLP classifier. It’s worth noting that MIOSTONE leverages optimized implementation techniques to significantly enhance training efficiency (Fig. 3**b** and Sec. 4.5). Most tree-agnostic models, provided by highly optimized toolkits (*e.g.,* Scikit-learn), remain the most efficient models. However, their training times increase much more rapidly than those of tree-aware methods as the number of samples grows. While tree-aware methods generally have slower training times compared to tree-agnostic methods, the relatively small size of microbiome datasets ensures that even the slowest tree-aware methods require only minutes to train. Given its enhanced predictive performance and model interpretability (Sec. 2.5), the computational cost will not hinder the applicability of MIOSTONE.

**Fig. 3.**
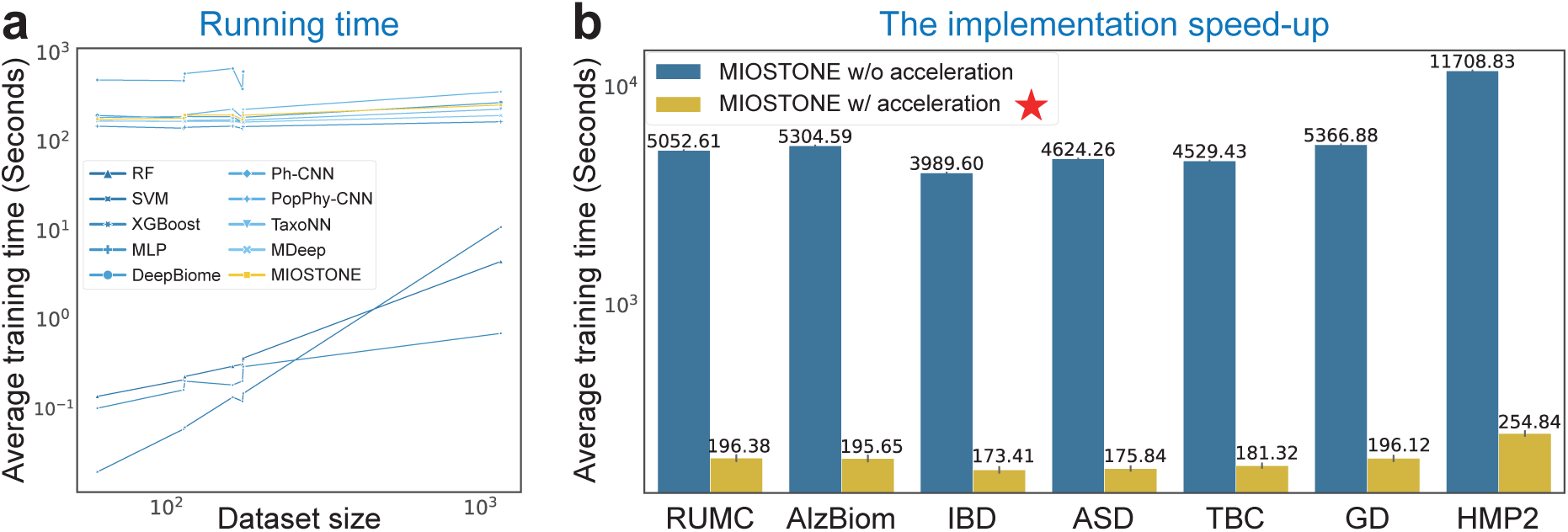
Runtime analysis. (**a**) MIOSTONE demonstrates comparable training efficiency to other tree-aware methods. Tree-agnostic methods, with their highly optimized implementations, are the most efficient models for small microbiome datasets. However, their training times escalate significantly faster than those of tree-aware methods as the sample size increases. (**b**) MIOSTONE utilizes a fully connected DNN with additional pruning, offering significantly greater efficiency than naive models based on customized taxonomic connections. The setting used by MIOSTONE is marked by ★.

MIOSTONE’s design incorporates several key components, including its taxonomy-encoding DNN architecture, data-driven aggregation of neuron representations, and taxonomy-dependent internal neuron dimensionality. Next, To assess the impact of each component on disease status prediction, we conducted several control studies in which we modified MIOSTONE by replacing its components with alternative solutions. Specifically, we considered multiple variants of MIOSTONE: (1) Replacing the GTDB-based taxonomy with an alternative taxonomy to encode the DNN architecture; (2) Replacing the taxonomy-encoding DNN architecture with a phylogeny-encoding alternative; (3) Making the data-driven aggregation of neuron representations deterministic by disabling stochastic gating; and (4) Setting the taxonomy-dependent internal neuron dimensionality to a fixed dimension of 2. For each variant, we applied MIOSTONE to seven real microbiome datasets with the same settings.

The results indicate that all key components positively and robustly contribute to MIOSTONE’s performance. In the first study (Fig. 4**a** and Fig. A.12), two variations of MIOSTONE equipped with two different taxonomic trees, exhibit comparable predictive performance across seven real microbiome datasets. This suggests that variations and inconsistencies between different taxonomic trees do not undermine the effectiveness of MIOSTONE. In the second study (Fig. 4**b** and Fig. A.13), we replaced the taxonomy with the phylogeny from the Web of Life (WoL) v2 database (Zhu et al, 2019). The WoL phylogeny is a dichotomous structure with varying depths across different branches, in contrast to the eight-layer structure of the taxonomy. The phylogeny-encoding architecture mirrors the dichotomous structure by enforcing connections between neurons based on their lineages. while MIOSTONE can emulate any hierarchical correlation among taxa within its architecture, alternatives such as phylogeny- encoding DNN architecture perform significantly worse than the taxonomy-encoding one. This suggests that phylogeny, as a more detailed scaffold for microbial classification, may present additional challenges during training compared to taxonomy. In the third study (Fig. 4**c** and Fig. A.11), the data-driven aggregation of neuron representations outperforms or matches the deterministic selection of nonlinear representation across all seven microbiome datasets. This underscores MIOSTONE’s key novelty in discerning whether taxa within the corresponding taxonomic group provide a more effective explanation of the trait when evaluated holistically as a group or individually as distinct taxa. In the last study (Fig. 4**d** and Fig. A.9), the taxonomy-dependent internal neuron dimensionality consistently outperforms the fixed dimensionality approach across all seven microbiome datasets, irrespective of sample sizes and feature dimensionality. This suggests that MIOSTONE’s assigning larger taxonomic groups with greater representation dimensionality can aid in capturing more complex biological patterns to predict traits, compared to using fixed representation dimensionality. Furthermore, MIOSTONE demonstrates robustness in selecting the hyperparameter *α* that controls the shrinkage of taxonomy-dependent representation dimensionality (Fig. 4**e** and Fig. A.8).

**Fig. 4.**
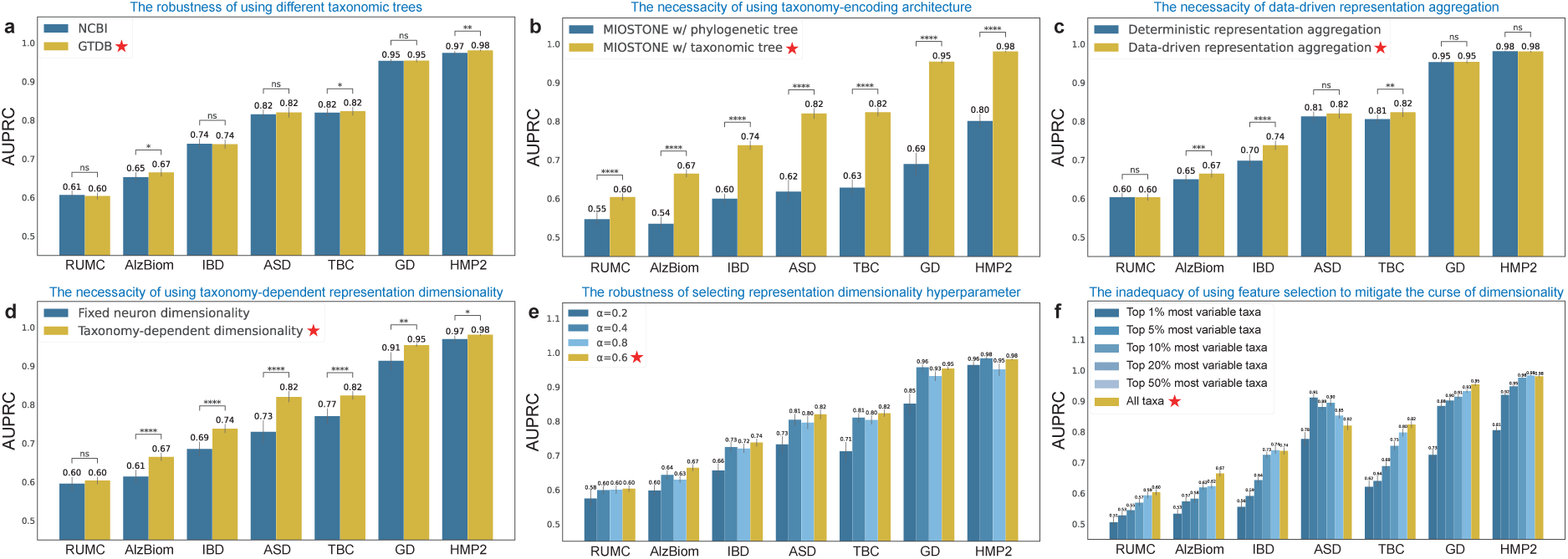
Dissecting the performance of MIOSTONE through control studies. (**a**) MIOSTONE demonstrates robustness across various taxonomic trees. Two variations of MIOSTONE utilizing taxonomies from GTDB and NCBI respectively, demonstrate comparable predictive performance across seven real microbiome datasets. (**b**) While MIOSTONE can emulate any hierarchical correlation among taxa within its architecture, alternatives, such as phylogenic trees, perform significantly worse than the taxonomy-encoding architectures. (**c**) MIOSTONE’s data-driven aggregation of neuron repre- sentations either outperforms or matches the performance of the deterministic selection of nonlinear representations across most datasets. (**d**) By assigning larger taxonomic groups greater representation dimensionality, MIOSTONE can capture more complex biological patterns for trait prediction, outperforming methods that use fixed representation dimensionality. (**e**) MIOSTONE demonstrates robustness in selecting the hyperparameter that controls taxonomy-dependent represen- tation dimensionality. (**f**) The curse of dimensionality cannot simply be mitigated using feature selection. MIOSTONE trained with all microbiome features, either outperforms or matches the performance of the model trained with a subset of highly variable taxa across most datasets. All settings used by MIOSTONE are marked by ★.

MIOSTONE’s superior performance stems from its taxonomy-encoding design, which effectively reduces model complexity and alleviates the curse of dimensionality (Liu et al, 2017). As a final step, we conducted an exploratory study to evaluate alternative strategies for encoding taxonomy and mitigating the curse of dimensionality. One alternative approach to encode taxonomy involves using the internal taxonomic units as additional features. Specifically, tree-agnostic models take input as concatenated features from both the taxa and the internal taxonomic units. Our analysis shows that Treating the internal taxonomic units as additional features results in marginal or even worse predictive performance, as measured by both AUPRC and AUROC (Fig. A.7). Furthermore, One may naturally wonder if we can mitigate the curse of dimensionality by only utilizing a subset of highly informative features. To answer this question, we chose the top-*k* highly variable taxa that contribute strongly to sample-to-sample variation, where *k* ranges from 1%, 5%, 10%, 20%, 50%, and 100% among all taxa. The selection of highly variable taxa is inspired by the identification of highly variable genes (Stuart et al, 2019) in single-cell RNA-seq, as implemented by the Scanpy python package (Wolf et al, 2018). For each taxa subset, we trained and evaluated MIOSTONE on seven real microbiome datasets using the same settings as for the full set. We found that MIOSTONE trained with all microbiome features either outperforms or matches the performance of the one trained with a subset of highly variable taxa on most of the datasets, with ASD being the only exceptions (Fig. 4**f** and Fig. A.10). It is crucial to note that the ASD dataset contains only 60 samples, profiled with 7,287 taxa. In such an extreme case of low sample size, we reasoned that proper feature selection tends to be beneficial in mitigating potential overfitting problems. Exploring the integration of feature selection into MIOSTONE could be an intriguing avenue for future research.

### 2.3 MIOSTONE improves predictive performance in sample-limited tasks through knowledge transfer

In microbiome data analysis, the transfer learning paradigm (Weiss et al, 2016) can be highly beneficial. This approach proves particularly beneficial for prediction tasks involving small datasets, as leveraging knowledge accumulated from existing models can substantially enhance predictive performance. To evaluate MIOSTONE’s ability to transfer knowledge from existing models, we selected two datasets (HMP2 and IBD) with the common objective of exploring the relationship between the gut microbiome and two inflammatory bowel disease subtypes: Crohn’s disease (CD) and ulcerative colitis (UC). We then investigated whether the knowledge acquired from the large HMP2 dataset with 1,158 samples could enhance the predictive performance on the smaller IBD dataset with 174 samples.

We pre-trained a model on the large HMP2 dataset and then utilized it for the smaller IBD dataset prediction task in three settings: (1) Directly employing the pre-trained HMP2 model for predictions on the IBD dataset (zero-shot); (2) Initializing the IBD model with the pre-trained HMP2 model and fine-tuning it on the IBD dataset (fine-tuning); and (3) Training a new model from scratch using only the IBD dataset (train from scratch). It’s worth noting that the HMP2 dataset has a higher feature dimensionality (10,614) than the IBD dataset (5,287). To ensure compatibility with the pre-trained model, we truncated the HMP features to match the dimensionality of the IBD dataset. It would be interesting to explore the direct use of the pre-trained, incompatible model in future research.

We then evaluated MIOSTONE’s performance using knowledge from pre-trained models, in terms of AUPRC (Fig. 5**a**) and AUROC scores (Fig. A.3**a**). Our analysis demonstrates that fine-tuning enhances predictive performance in MIOSTONE compared to training from scratch. For scientific rigor, the performance between fine-tuning and training from scratch is quantified using one-tailed two-sample t-tests to calculate p-values. These p-values confirm that the performance superiority of fine-tuning over training from scratch is both statistically significant and qualitatively discernible. In contrast, the other tree-aware baseline methods demonstrated varying degrees of success in transferring knowledge from existing models. For example, similar to MIOSTONE, TaxoNN showed enhanced predictive performance, though it still did not surpass MIOSTONE’s performance. However, MLP and MDeep did not exhibit a noticeable improvement with the pre-trained model, suggesting that the knowledge learned by the pre-trained model was completely overwritten by fine-tuning. Furthermore, PopPhy-CNN exhibited a reverse effect, experiencing a 15.2% decline in AUPRC and a 17.9% decline in AUROC scores with fine-tuning compared to training from scratch. This suggests that the knowledge learned by the pre-trained model may even hinder learning from the new data.

**Fig. 5.**
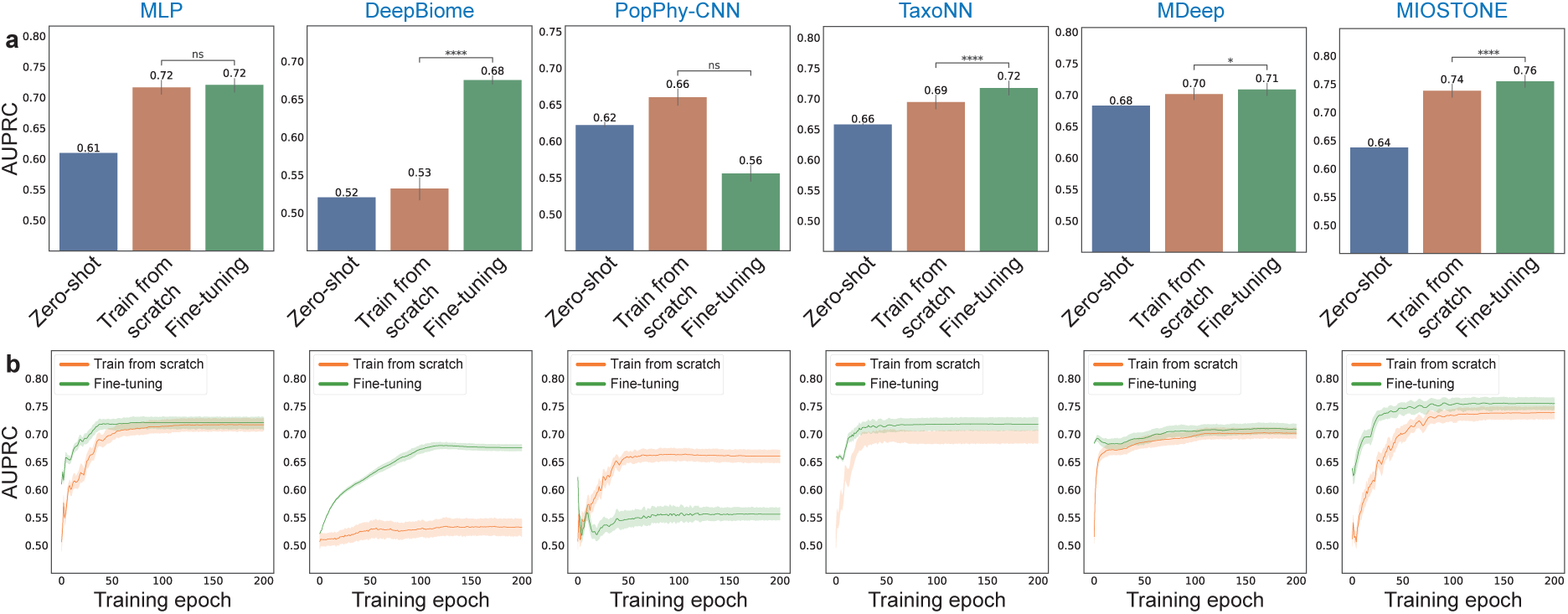
MIOSTONE enhances disease prediction by transferring knowledge from pre-trained models. (**a**) A model on the large HMP2 dataset is pre-trained and then employed for the smaller IBD dataset in three settings: direct prediction on IBD (*i.e.,* zero-shot), fine-tuning on IBD, and training IBD from scratch. Only tree-aware methods and MLP are included in the comparison, as most tree-agnostic methods are not well-suited for fine-tuning. Among the tree-aware methods, Ph-CNN is excluded because it is not scalable for processing the large HMP2 dataset. The prediction is conducted across three settings 20 times with varied train-test splits, and reported by the average performance assessed by AUPRC, along with 95% confidence intervals. (**b**) The training dynamics of various models were evaluated by comparing fine-tuning with training from scratch, analyzing AUPRC on test splits across different training epochs. MIOSTONE’s fine-tuning achieved better performance than training from scratch, requiring fewer training epochs.

Finally, we examined the MIOSTONE training dynamics by comparing fine-tuning with training from scratch, focusing on the reported performance on test splits across different training epochs (Fig. 5**b** and Fig. A.3**b**). We observed that MIOSTONE’s fine-tuning reaches a plateau and attains optimal performance within 40 out of 200 epochs, requiring significantly fewer training epochs for superior performance compared to training from scratch. In other words, leveraging knowledge from pre-trained models through fine-tuning empowers MIOSTONE. Since MIOSTONE is already computationally efficient, this approach can further reduce training time. We conclude that MIOSTONE effectively improves disease prediction through knowledge transfer via fine-tuning.

### 2.4 MIOSTONE learns meaningful and discriminative representations

While MIOSTONE was primarily developed for prediction, its exceptional performance in distinguishing disease status suggests that understanding the model’s internal mechanisms could provide valuable insights for scientific discovery. To this end, we began by investigating whether internal neuron representations within the MIOSTONE model encode disease-specific signatures. Representation learning is renowned for extracting meaningful high-level semantics from raw data and has been widely employed to uncover hidden patterns in biological and biomedical data (Bengio et al, 2013; Iuchi et al, 2021).

We extracted MIOSTONE’s internal neuron representations from three taxonomic levels (species, genus, and family), comparing them against other tree-aware methods. Given the drastically different DNN architectures of these methods, we extracted the last-layer latent representations, which are believed to encode the maximum semantic information, for a fair comparison of these tree-aware methods. We projected the extracted representations from all methods, which encode the semantic meanings of the input samples, into a two-dimensional embedding space using Principal Component Analysis (PCA). We then evaluated the effectiveness of these representations in distinguishing different disease statuses.

The patient samples with different disease statuses cannot be distinguished initially, considering that microbiome data is typically high-dimensional and noisy. For example, the PCA visualization of microbiome features based on taxa profiling fails to distinguish IBD disease subtypes, as samples from Crohn’s disease (CD) and ulcerative colitis (UC) patients are mixed together (Fig. 6). The other tree-aware methods demonstrated varying degrees of improvement in distinguishing between the two IBD disease subtypes. For example, models like MLP, DeepBiome, and TaxoNN struggled to differentiate between IBD disease subtypes, while MDeep demonstrated a clear separation between these subtypes. However, when we represented each patient sample by MIOSTONE’s internal neuron representations from even the bottom taxonomic level with least encoded semantics (*i.e.,* species), the resulting representations exhibited significantly improved separation among disease subtypes, suggesting that the model’s internal representations effectively capture diverse disease-specific signatures. The separation between disease subtypes is quantitatively measured by the silhouette value (Rousseeuw, 1987). MIOSTONE’s internal neuron representations exhibit higher silhouette values, suggesting greater similarity of each sample to its own disease subtype compared to other subtypes.

**Fig. 6.**
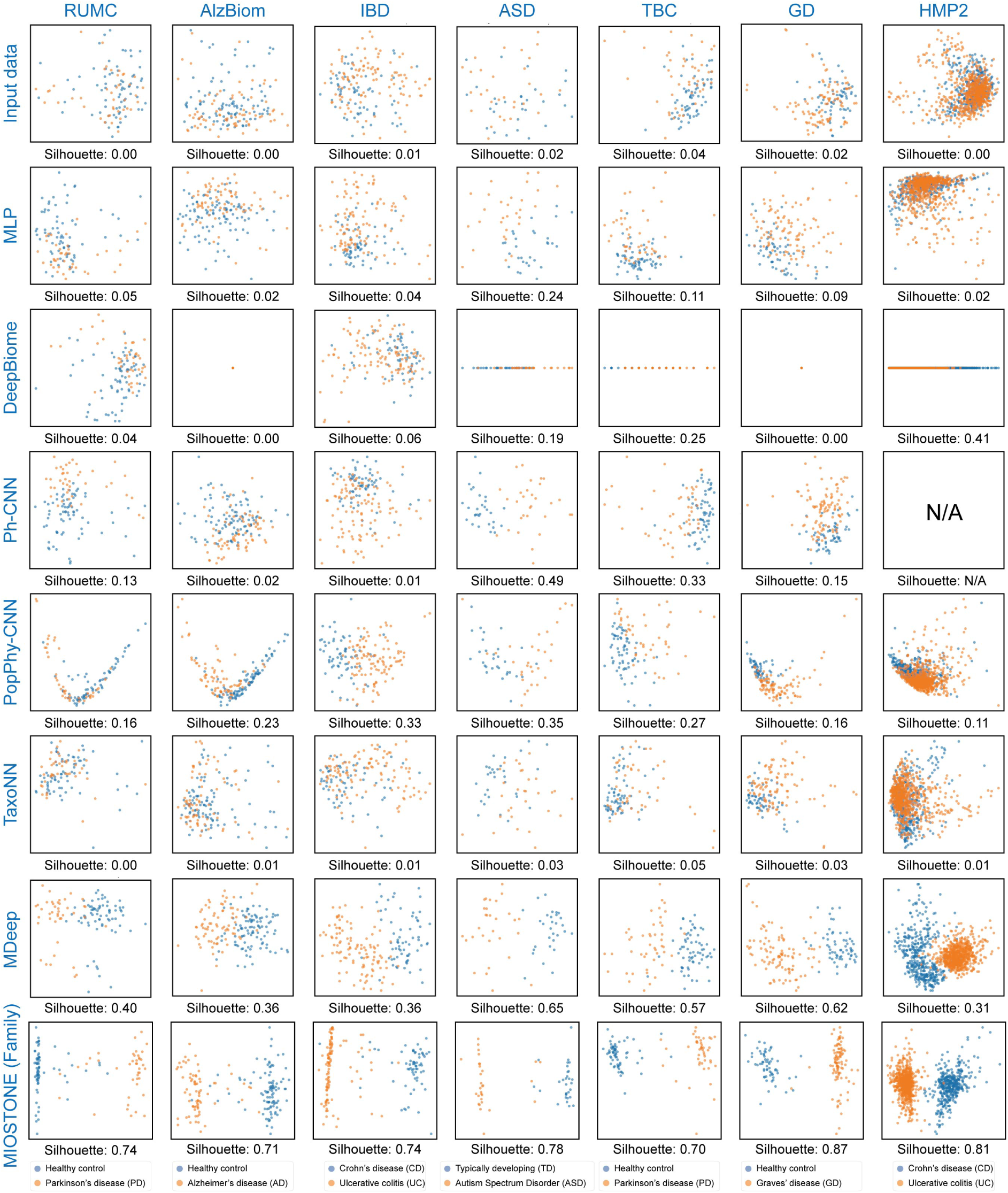
MIOSTONE learns meaningful and discriminative representations. MIOSTONE’s internal neuron representations of samples are projected onto a two-dimensional Principal Component space and evaluated their efficacy in distinguishing between different disease subtypes. MIOSTONE’s family level representations are compared to the last-layer latent representations of other tree-aware methods, with the exception of Ph-CNN on the HMP2 dataset, as Ph-CNN is not scalable for that dataset. MIOSTONE’s representations show significantly improved separation between disease subtypes, indicating that the model’s internal representations effectively capture diverse disease-specific signatures. This separation is quantitatively assessed using the silhouette value.

Finally, one may wonder whether the representations are independent of the data and purely a result of the model’s taxonomy-encoding architecture (Adebayo et al, 2018). If that were the case, drawing reliable and convincing conclusions from the model would be unwarranted. To address this, we conducted a sanity check by using an untrained MIOSTONE model to project the internal neuron representations of samples into a two-dimensional Principal Component space. We then evaluated their ability to distinguish between different disease subtypes and compared these results with those from the trained MIOSTONE model. We observed that the untrained model showed no separation between disease subtypes, confirming that MIOSTONE’s internal representations are data-dependent, effectively capturing disease-specific signatures during training (Fig. A.4).

### 2.5 MIOSTONE identifies microbiome-disease associations with high interpretability

Recognizing the model’s potential in capturing disease-specific semantics, we further delved into the MIOSTONE model to uncover significant microbiome-disease associations. Important associations were scored using feature attribution methods, which assign importance scores to taxonomic groups, with higher scores indicating greater importance to the model’s prediction. In this study, we employed three representative feature attribution methods, DeepLIFT, integrated gradient, and SHAP, to elucidate the relationship between microbiome taxa and disease trait without assuming any specific model architecture. We discovered that different feature attribution methods demonstrate strong consistency in quantifying crucial microbiome-disease associations from the trained MIOSTONE model (Fig. 7**a**). Furthermore, feature attribution methods demonstrated strong consistency in quantifying crucial microbiome-disease associations across different data splits (Fig. A.5**b**). Therefore, the consistency among different feature attribution methods and robustness across various data splits justify presenting only the DeepLIFT results hereafter. Since each feature attribution method evaluates the sample-wise contribution of each taxa within the sample to predicting the corresponding disease status, we calculated the overall importance of each taxa by aggregating its contributions across all samples. For robustness, we repeat this procedure for each data split across 20 repetitions and aggregated the scores into a consensus importance score for interpretation.

**Fig. 7.**
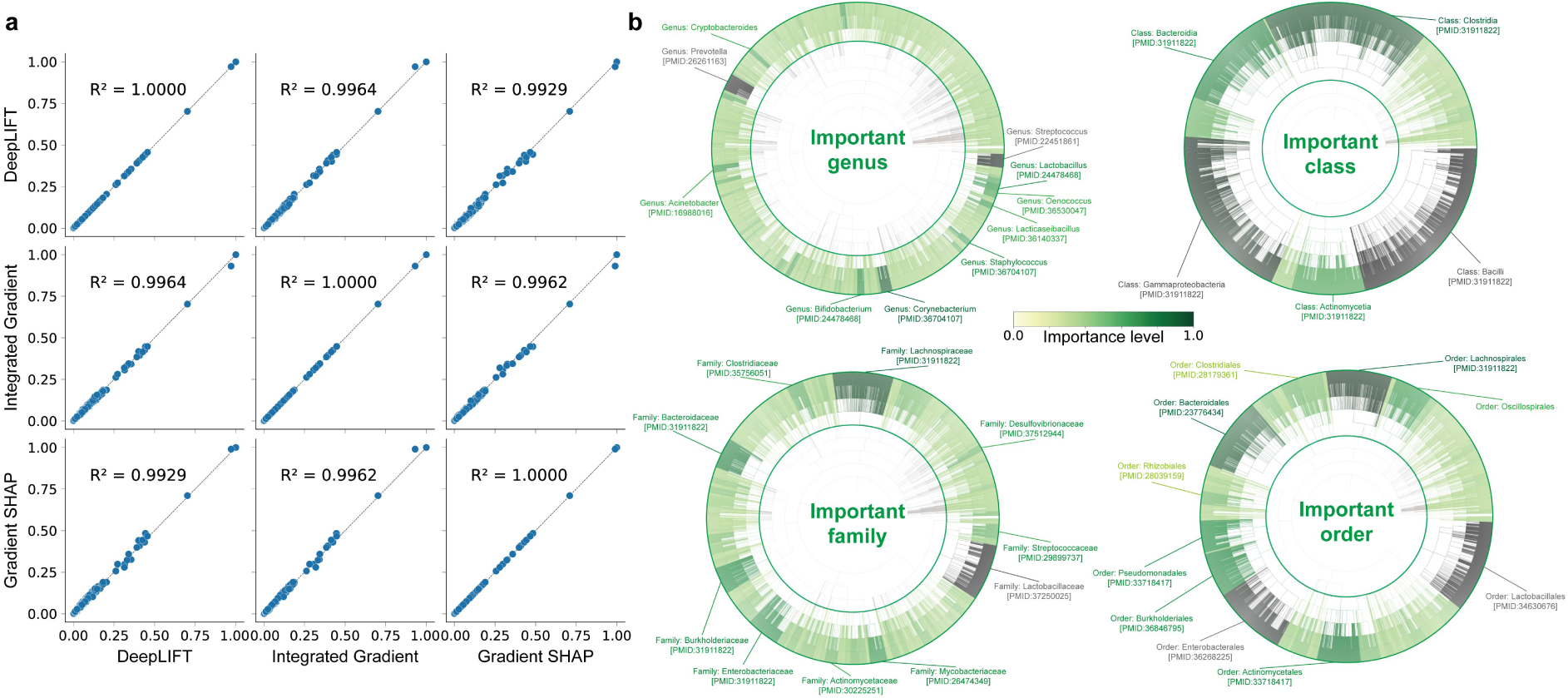
MIOSTONE discovers microbiome-disease associations across different taxonomic levels. Feature attribution methods are used to interpret the MIOSTONE model and quantify the relationships between microbiome taxa and disease traits. (**a**) Three mainstream feature attribution methods–DeepLIFT, Integrated Gradients, and SHAP– demonstrate strong consistency in quantifying key microbiome-disease associations derived from the MIOSTONE model. (**b**) Feature importance attribution by DeepLIFT is performed at the genus, family, order, and class taxonomic levels. When emphasizing an important taxonomic group, the taxonomic subtree rooted at that group—along with its group-specific taxa—is highlighted for improved visualization. The top-ranked taxonomic groups are displayed with their respective names, supported by relevant literature evidence and accompanying PubMed identifiers.

We initially focused on identifying important microbiome-disease associations in differentiating between two IBD disease subtypes: Crohn’s disease (CD) and ulcerative colitis (UC), at the genus level. We highlighted top-ranked genera reported by DeepLIFT with their respective names, supported by literature evidence with accompanying PubMed identifiers (Fig. 7**b**). For example, the microbial community associated with *Prevotella* has been reported to have significantly different abundance in UC compared to controls as well as CD (Kabeerdoss et al, 2015). Furthermore, studies have reported that the detection frequency of *Streptococcus* in UC patients was significantly higher than in healthy subjects. Infection with highly virulent specific types of Streptococcus might be a potential risk factor in the aggravation of UC (Kojima et al, 2012).

The identification of significant microbiome-disease associations in distinguishing between IBD disease subtypes can be further extended to coarser resolutions such as family-level, order-level, and class-level. For example, the *Lachnospiraceae* family, predominantly found in the gut microbiota of mammals and humans, has been reported to have significantly different abundance between IBD disease and health controls (Lee et al, 2020; Sasaki et al, 2019). Moreover, studies have reported increased levels of the *Bacilli* and *Clostridia* classes in UC patients, while the levels of *Clostridia* and *Bacteroidia* are decreased in CD patients (Alam et al, 2020).

One may wonder whether the microbiome-disease associations reported by DeepLIFT are independent of the data and purely a result of the model’s taxonomy-encoding architecture (Adebayo et al, 2018). If that were the case, drawing reliable and convincing microbiome-disease associations using feature attribution methods would be unwarranted. As a sanity check, we used an untrained MIOSTONE model and evaluated the consistency between the microbiome-disease associations it reported and those from a trained model (Fig. A.5**c**). The low consistency suggests that the microbiome-disease associations reported by feature attribution methods are data-dependent. We conclude that MIOSTONE effectively identifies microbiome-disease associations across different taxonomic levels, providing valuable insights for scientific discovery.

## 3 Discussion

In this study, we propose MIOSTONE, an accurate and interpretable machine learning method for investigating microbiome-trait associations. At its core, MIOSTONE leverages the intercorrelation of microbiome features based on their taxonomic relationships. The key novelties of MIOSTONE are three-fold: (1) the taxonomy-encoding architecture harnesses the capabilities of DNNs with mitigated concerns of overfitting; (2) the ability to determine whether taxa within the corresponding taxonomic group provide a better explanation in a data-driven manner; and (3) the interpretable architecture facilitates the understanding of microbiome-trait associations. We validated its performance on seven real datasets, demonstrating its superiority in predictive performance and biological interpretability. Beyond disease status prediction, it can discover significant microbiome-disease associations and transfer knowledge to enhance predictive performance in tasks with limited samples.

Methodologically, our approach provides a systematic way to circumvent the fundamental computational challenges in the conventional analysis of microbiome data. First of all, the curse of dimensionality has long been a dilemma in computational modeling. Specifically, the relatively low number of samples may lead to overfitting during training, thereby impeding the use of powerful analysis tools like DNNs at the expense of prediction accuracy. The hierarchical neural network framework adopted by MIOSTONE significantly reduces the model’s complexity, thereby mitigating the curse of dimensionality and overfitting. Moreover, the biologically-informed neural networks are highly generic and expressive, allowing for the representation of any hierarchical relationships or functional dependencies among microbial taxa. By incorporating this knowledge into biologically-informed neural networks, we can attribute the information encoded by the data to these pre-specified biological concepts, offering a natural interpretation of the model’s internal mechanisms.

Numerous studies have focused on accurately differentiating disease states and understanding the differences in microbiome profiles between healthy and ill individuals. Most of them primarily focus on various statistical methods, without explicitly modeling the underlying molecular mechanisms that give rise to nonlinearity and microbe-microbe interactions among a large number of microbial taxa, which in principle drive microbiome dynamics. We hypothesized that this might be due to the fact that the curse of dimensionality already makes first-order association identification highly challenging, let alone the detection of higher-order interactions, which is a much more difficult task. Given that the curse of dimensionality has been systematically mitigated, a potential research direction for future studies is to quantify important higher-order microbe-microbe interactions instead of focusing solely on an individual taxon. This could involve using feature interaction detection methods developed in the interpretable machine learning community (Tsang et al, 2018; Chen et al, 2024b,a).

Furthermore, to the best of our knowledge, MIOSTONE is the first attempt to successfully introduce the transfer learning paradigm into microbiome data analysis and systematically evaluate how leveraging knowledge from existing models can substantially enhance predictive performance. This is because previous works either rely on tree-agnostic models, which are not suitable for transfer learning, or tree-aware models, which are not powerful enough to model the data effectively. In this study, we demonstrated the transfer learning paradigm using the IBD and HMP2 datasets–the only publicly accessible dataset pair that shares the same disease type but differs significantly in sample size. It is important to note that MIOSTONE is just the beginning of efforts to build powerful models that extract valuable insights from small microbiome datasets to enhance new analyses.

Lastly, the evaluation of detected important microbiome-disease associations or microbe-microbe interactions relies solely on literature support. While this approach is reasonable for evaluation purposes, it might limit the credibility for less studied taxa. A potential research direction for future studies is to provide confidence estimation for the top-ranked microbiome-disease associations to complement the literature support, using measures such as q-values (Storey, 2003), with the assistance of the recently proposed knockoffs framework (Barber and Candèes, 2015; Lu et al, 2018).

In conclusion, MIOSTONE adeptly navigates the analysis of microbiome data, effectively addressing issues such as data imperfections, sparsity, low signal-to-noise ratio, and the curse of dimensionality. We believe that this powerful analytical tool will enhance our understanding of the microbiome’s impact on human health and disease and will be instrumental in advancing novel microbiome-based therapeutics.

## 4 Methods

### 4.1 Datasets

We conducted a comprehensive evaluation of MIOSTONE using seven publicly available microbiome datasets with varying sample sizes and feature dimensionality (Details in Tab. A.1). The AlzBiom dataset (Laske et al, 2022) explored the relationship between the gut microbiome and Alzheimer’s disease (AD). It comprises 75 amyloid-positive AD samples and 100 cognitively healthy control samples from the AlzBiom study, profiled with 8,350 taxa. The ASD dataset (Dan et al, 2020) investigated the connection between the gut microbiome and abnormal metabolic activity in Autism Spectrum Disorder (ASD). It comprises 30 typically developing (TD) and 30 constipated ASD (C-ASD) samples, profiled with 7,287 taxa. The GD dataset (Zhu et al, 2021) explored the relationship between the gut microbiome and Graves’ disease (GD). It comprises 100 GD samples and 62 healthy control samples, profiled with 8,487 taxa. The TBC and RUMC datasets (Boktor et al, 2023) are two cohort studies investigating the connection between the gut microbiome and the Parkinson’s disease (PD). The TBC cohort includes 46 PD samples and 67 healthy control samples, profiled with 6,227 taxa. The RUMC cohort comprises 42 PD samples and 72 healthy control samples, profiled with 7,256 taxa. The IBD dataset (Gonzalez et al, 2022) investigated the relationship between the gut microbiome with two primary subtypes of inflammatory bowel disease (IBD): Crohn’s disease (CD) and ulcerative colitis (UC). It comprises 108 CD and 66 UC samples, profiled with 5,287 taxa. The HMP2 dataset (Lloyd-Price et al, 2019) in the Integrative Human Microbiome Project (iHMP) also investigated the relationship between the gut microbiome and two IBD subtypes: CD and UC. Compared to the IBD dataset, this dataset expands the sample size to 1,158 (728 CD and 430 UC samples), with an expanded taxa set of size 10,614.

Additionally, we created three simulated datasets representing various settings. Two simulated datasets were generated by using the microbiome data simulator MIDASim (He et al, 2024). We used a microbiome data simulator MIDASim (He et al, 2024) to generate simulated data by leveraging a template microbiome dataset while preserving its correlation structure to maintain similarity. We selected the IBD dataset as the template and adjusted the simulation settings to create two datasets with varying levels of difficulty in distinguishing between labels. Since MIDASim is not designed to generate simulated samples with labels, we selected a real dataset with positive and negative labels. We then simulated we generated the same number of positive and negative samples as the IBD dataset before combining them into a unified simulated dataset. The third simulated dataset was generated by Zhai et al (2024) with a small sample size and low feature dimensionality to evaluate MIOSTONE’s performance in an atypical setting. The dataset comprises 48 genus and 50 samples, comprising 32 positive and 18 negative samples, respectively. (Refer to Sec. A.1 for the details of simulated dataset generation).

### 4.2 Taxonomic tree

The abundance features in these datasets were profiled using the standard operating procedure for shotgun metagenomics implemented in the Qiita platform (Gonzalez et al, 2018). Specifically, the raw sequencing reads were processed using fastp and Minimap2 to remove low-quality sequences, adapter sequences and sequences that are susceptible to host contamination (Armstrong et al, 2022). The processed reads were classified using Woltka v0.1.4 (Zhu et al, 2022) against the Web of Life (WoL) v2 database (Zhu et al, 2019), which includes 15,953 microbial genomes. WoL offers taxonomic annotations, which are based on the Genome Taxonomy Database (GTDB) (Parks et al, 2018), for these microbial genomes, spanning 124 phyla, 320 classes, 914 orders, 2,057 families, 6,811 genera, and 12,258 species. Given that each dataset only profiles a subset of taxa within the WoL taxonomy, the MIOSTONE model is constructed using a pruned taxonomy tree tailored for each dataset.

The pruning process retains only the taxa shared between the WoL taxonomy and those profiled in a given dataset. Since the train/test split uses the same subset of taxa as features, taxonomy pruning does not result in any data leakage. It is important to note that pruning the taxonomy tree does not impact predictive performance because non-shared features, even if retained, are effectively treated as zero through imputation. (Refer to Fig. 4 and Fig. A.12 for the robustness of MIOSTONE across various sources of taxonomic trees).

### 4.3 Baseline methods

We evaluate the performance of MIOSTONE in comparison to nine baseline methods, divided into two categories: tree-agnostic methods and tree-aware methods. The former category comprises random forest (RF), support vector machine (SVM) with a linear kernel, XGBoost (Chen and Guestrin, 2016), and multi-layer perceptron (MLP), while the latter includes DeepBiome (Zhai et al, 2024), Ph-CNN (Fioravanti et al, 2018), PopPhy-CNN (Reiman et al, 2020), TaxoNN (Sharma et al, 2020), and MDeep (Wang et al, 2021).

We used the Scikit-learn implementation (Pedregosa et al, 2011) with default settings for the RF and SVM models, and the official implementation for the XGBoost classifier. For the MLP classifier, we configured a pyramid-shaped architecture with one hidden layer of size half of the input dimensionality. For other tree-aware models, we used the recommended implementation settings. (Refer to Sec. A.2 for the details of model settings and training). Unless explicitly specified in Sec. A.2, we preprocessed the microbiome features using centered log-ratio transformation (Aitchison, 1982).

### 4.4 MIOSTONE design

MIOSTONE trains a deep neural network to predict disease traits from microbial taxa abundance profiles, with its architecture that precisely mirrors the taxonomic hierarchy based on the Genome Taxonomy Database (GTDB) (Parks et al, 2018). Each neuron in the network corresponds to a specific taxonomic group, and the connections between neurons represent the subordination relationships between these groups, such as “species A belongs to genus B” or “genus B belongs to family C” relationships. Unlike fully connected DNNs, MIOSTONE only connects neurons that are directly related in the taxonomic hierarchy. This design choice significantly reduces the model’s complexity, effectively mitigating the overfitting problem, while simultaneously enhancing its interpretability.

We denote our input training data set as *D* = {(*x*_1_*, y*_1_),(*x*_2_*, y*_2_),· · ·,(*x_n_, y_n_*)}, where *n* is the number of samples. For each sample *i*, *x_i_* ∈ R*^p^* represents the *p*-dimensional profiled abundance of microbial taxa as features, and *y_i_*∈ R denotes the corresponding trait label, which can be either binary (*e.g.,* disease status) or continuous (*e.g.,* age). Our goal is to learn a predictive function R*^p^* 1→ R, parameterized by a DNN, that accurately predict the trait label *y* ∈ R for the microbiome sample *x* ∈ R*^p^*.

One challenge in modeling microbiome-trait associations is the ambiguity in fragment-to-taxon assignments. For example, a sequenced viral fragment from the Omicron variant may be mistakenly assigned to the Delta variant, both belonging to the SARS-CoV-2 lineage, but it is unlikely to be assigned to SARS-CoV-1. To tackle this issue, MIOSTONE employs a data-driven strategy to determine whether taxa within the corresponding taxonomic group provide a better explanation for the disease traits when considered holistically or individually. This strategy aims to balance the reduction of ambiguity in fragment-to-taxon assignments with the effective explanation of the trait of interest. The underlying rationale is that each taxonomic group may exhibit different levels of ambiguity in these assignments, necessitating distinct treatment. High ambiguity suggests that taxa within a group may not be individually meaningful and should be considered collectively, while low ambiguity implies that each taxon may have an individual impact on the trait of interest.

We implement this strategy by introducing a stochastic gate (Louizos et al, 2018) for each internal neuron (Fig. 1**b**). Specifically, an internal neuron *v* is characterized by two multi-dimensional representations: an additive representation 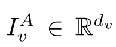 and a nonlinear representation 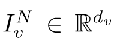, where *d_v_* is the representation dimension. The additive representation 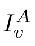 is obtained by concatenating the additive representations of all children of *v* and then applying a linear transformation:

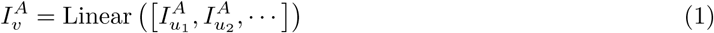

where *u*_1_*, u*_2_,· · · are the children of *v* and Linear(·) is a linear transformation function. To obtain the nonlinear representation 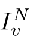, we first concatenate the nonlinear representations of all children of *v* and then apply a multi-layer perceptron (MLP) to transform it into an intermediate non-linear representation:

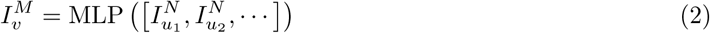

During the training phase, the additive representation and the nonlinear representation compete against each other for improved trait prediction through a stochastic gate *m_v_*∈ (0, 1) that combines 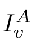 and 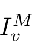 into the final nonlinear representation 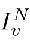 :

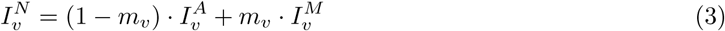

where the gate *m_v_*is based on the hard concrete distribution (Louizos et al, 2018), which is a differentiable relaxation of the Bernoulli distribution:

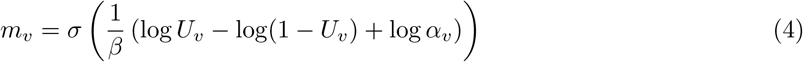

where 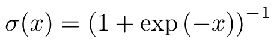 is the sigmoid function and *U_v_*∼ Uniform(0,1) an independent random variable following a continuous uniform distribution. This relaxation is parameterized by a trainable parameter *α_v_* and a temperature coefficient *β* ∈ (0,1) controlling the degree of approximation. As *β* → 0, the gate *m_v_* converge to a Bernoulli random variable. We set *β* = 0.3 in our experiments. When the gate *m_v_* has a value close to 1 (*i.e.,* in “on” state), all taxa within the group (*e.g., X*_1_ and *X*_2_ in Figure 1(B)) will be selected to contribute to the prediction individually. When the gate *m_v_*has a value close to 0 (*i.e.,* in “off” state), all taxa within the group may not be as individually meaningful as when considered holistically as a group.

Intuitively, larger taxonomic groups should possess a greater representation dimension to capture potentially more complex biological patterns. However, the dimension should not become excessively large, as this might lead each taxonomic group to merely memorize information from its descendants, rather than distill and learn new patterns. Thus, we determine the representation dimension *d_v_* for each internal neuron *v* recursively as 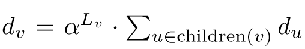 where children(*v*) denotes the children of *v*. Here, *α* is a hyperparameter controlling the shrinkage of the representation dimension and *L_v_*is the taxonomic level of *v*, starting from *L* = 1 for species and increasing by 1 for each level up to *L* = 7 for domains. We set *α* = 0.6 in our experiments.

Tracing the taxonomy tree up from the leaf nodes to the root, we obtain the nonlinear representations of the root node 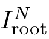 . We then apply a batch normalization layer (Ioffe and Szegedy, 2015) to 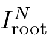, and feed it into a MLP classifier to predict the trait label *y*: Batch normalization (BN) (Ioffe and Szegedy, 2015) assists in mitigating the influence of internal covariate shift caused by different taxonomic groups. The training objective is to minimize the cross-entropy loss between the predicted label and the ground truth label:

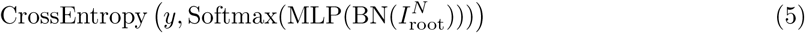

### 4.5 MIOSTONE implementation and training

The MIOSTONE model is designed to only connect neurons that are directly related in the taxonomic hierarchy. Unlike fully connected DNNs, which are efficiently optimized for parallel training by off-the-shelf deep learning libraries, the MIOSTONE model with customized hierarchical connections lacks similar optimization. In essence, pursuing interpretability in model architecture may come at the cost of computational efficiency. This is due to the need to calculate and sequentially backpropagate gradients with respect to model parameters layer-by-layer, without leveraging the speed-up provided by parallelism. To attain both interpretability and computational efficiency concurrently, we construct an equivalent MIOSTONE model by employing a fully connected DNN with additional pruning. Specifically, we initially construct a 7-layer fully connected DNN, with each layer corresponding to a specific taxonomic level. We subsequently utilize pruning techniques (Han et al, 2015) to remove connections between neurons that are not relevant in the taxonomic hierarchy, retaining only the taxonomy-encoding connections. This approach ensures that the pruned DNN not only maintains interpretability in terms of taxonomic hierarchy but is also efficiently optimized for parallel training by off-the-shelf deep learning libraries (Fig. 3**b**).

To train the MIOSTONE model, we initialize all weights uniformly at random. We optimize the objective function (Eq. 5) using ADAM with decoupled weight decay (Loshchilov and Hutter, 2019), a commonly-used stochastic gradient descent algorithm, with an initial learning rate of 0.001. Given the small size of microbiome datasets, we train the model for 200 epochs with a batch size of 512 to ensure convergence. We used 5-fold cross-validation, trained separate models on each training split and evaluated them on the corresponding test splits. We implemented MIOSTONE using the PyTorch library (Paszke et al, 2019) on NVIDIA RTX A6000 GPUs. (Refer to Sec. A.2 for more details).

### 4.6 Model interpretation

We evaluate the potential of MIOSTONE in uncovering significant microbiome-disease associations. We quantify important associations using feature attribution methods, which assign importance scores to taxonomic groups, with higher scores indicating greater importance to the model’s prediction. In this study, we employ three representative model-agnostic feature attribution methods to elucidate the relationship between microbiome taxa and disease trait without assuming any specific model architecture. In brief, the first method, DeepLIFT (Shrikumar et al, 2017), compares the activation of each neuron to its reference activation and assigns contribution scores based on the disparity. The second method, integrated gradient (Sundararajan et al, 2017), involves summing over the gradient with respect to different scaled versions of the input. And the third method, SHAP (Lundberg and Lee, 2017), explains the contribution of each feature to the prediction based on the game-theoretically optimal Shapley values. However, in principle, any off-the-shelf feature attribution methods can be utilized.

### 4.7 Code and data availability

The software implementation and all models described in this study are available at https://github.com/batmen-lab/MIOSTONE. All public datasets used for evaluating the models are available at https://github.com/batmen-lab/MIOSTONE/tree/main/data.

## Declarations

- The research is supported by the Canadian NSERC Discovery Grant RGPIN-03270-2023.
- The authors declare that they have no conflict of interest.
- There is no ethics approval and consent to participate involved in this study.
- There is no consent for publication involved in this study.
- Code and data availability: https://github.com/batmen-lab/MIOSTONE.
- Author contribution: Y.J. implemented the code and did the analysis. M.A. and Q.Z. set up and preprocessed the datasets. Y.L. and Y.J. prepared the figures. Y.L. wrote the main manuscript text. All authors participated the discussion. All authors reviewed the manuscript.

## A Appendix

**A.1 Dataset details**

MIOSTONE used seven publicly available microbiome datasets with varying sample sizes and feature dimensionality, with details listed in Tab. A.1 and Fig. A.1.

**Table A1.** The details of the real datasets investigated by MIOSTONE.

| Dataset | Sample size | Feature size | Data source | Notes |
| --- | --- | --- | --- | --- |
| AlzBiom | 175 samples (75 amyloid-positive and 100 healthy control) | 8,350 taxa | EBI-ENA ID: PRJEB47976 | The sequencing data is clean (QC'ed, host-filtered). |
| ASD | 60 samples (30 typically developing and 30 constipated ASD) | 7,287 taxa | EBI-ENA ID: PRJNA451479 | The sequencing data is raw (non-QC'ed). |
| GD | 162 samples (100 Graves' disease and 62 healthy control) | 8,487 taxa | EBI-ENA ID: PRJNA602729, PRJNA602731, PRJNA602732, PRJNA638403, PRJNA638404, PRJNA638405 | Most samples have a pair of FASTQ files. However, 4 samples (three.lst) have a third, unpaired FASTQ file that is very small, and it should be excluded from the analysis. 12 samples have only one FASTQ file, which appears to be single-end sequences. Two samples: GA61 (SRR12000211) and GA89 (SRR12005695) are missing from the metadata. Therefore they were dropped from the data. |
| RUMC | 114 samples (42 Parkinson's disease and 72 healthy control) | 7,256 taxa | Qiita ID: 12975 | 20 samples in BIOM are missing in metadata. These samples were dropped. |
| TBC | 113 samples (46 Parkinson's disease and 67 healthy control) | 6,227 taxa | Qiita ID: 14476 | 5 samples in BIOM are missing in metadata. These samples were dropped. |
| IBD | 174 samples (108 Crohn's disease and 66 ulcerative colitis) | 5,287 taxa | Qiita ID: 12675 | The dataset contains metagenomic sequencing data and associated metadata. More details can be found at: https://qiita.ucsd.edu/study/description/12675 |
| HMP2 | 1,158 samples (728 Crohn's disease and 430 ulcerative colitis) | 10,614 taxa | Qiita ID: 11484 | The dataset contains metagenomic sequencing data and associated metadata from the Human Microbiome Project. More details can be found at: https://hmpdacc.org/ihmp |

**Fig. A1.**
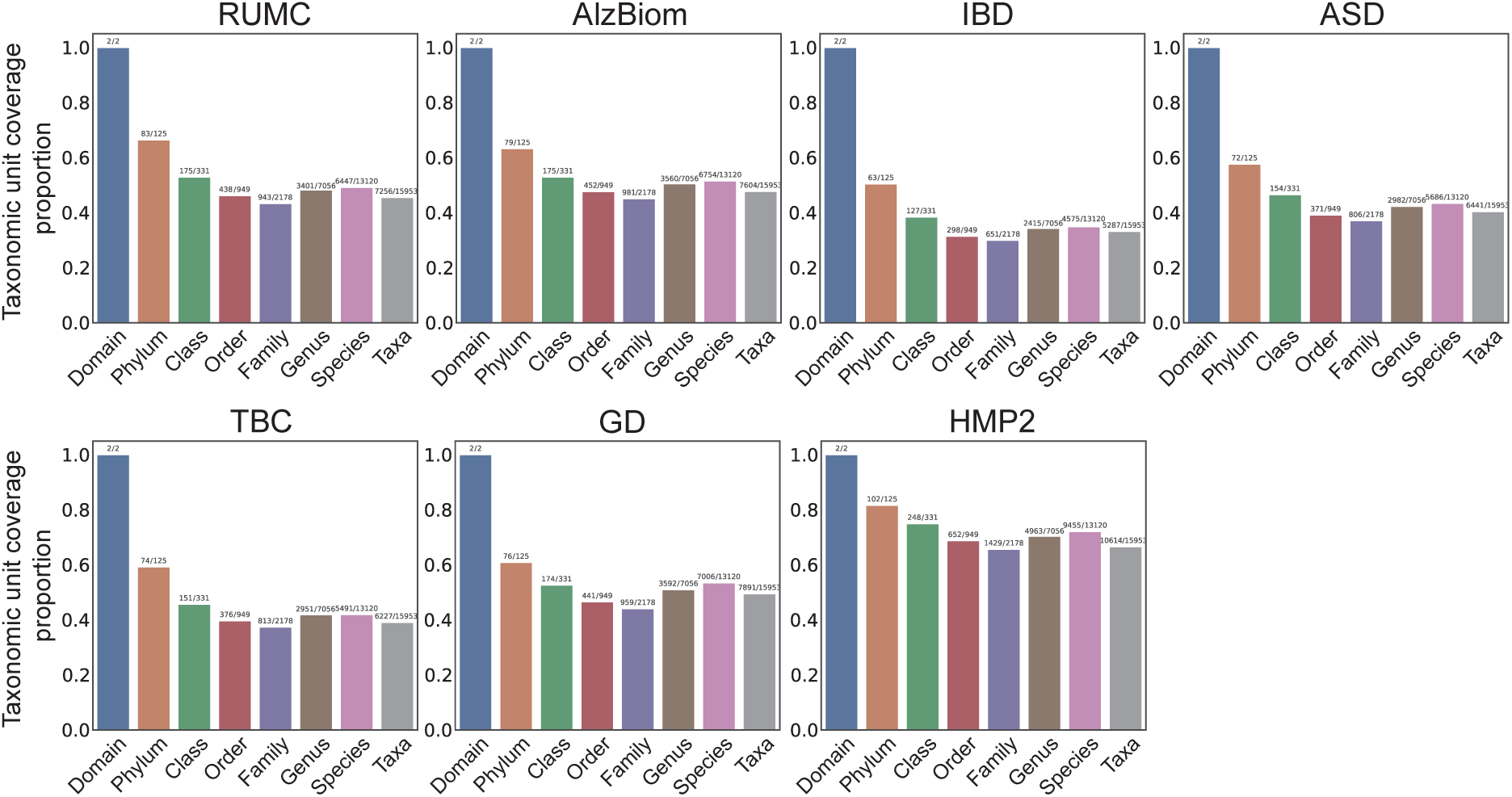
The microbiome datasets exhibit varying microbial taxa sizes and encompass different proportions of taxonomic levels

Additionally, we created two simulated datasets using the microbiome data simulator MIDASim (He et al, 2024). MIDASim generates simulated data by leveraging a template microbiome dataset and pre- serving its correlation structure to ensure similarity. Specifically, we employed the parametric mode of MIDASim, utilizing a generalized gamma distribution to model the relative abundances of microbiome data. This approach is tailored for simulation studies that involve modifying the log-mean relative abun- dance. Since MIDASim is not designed to generate simulated samples with labels, we selected a real dataset with positive and negative labels. We then simulated samples separately for each label (posi- tive and negative) before combining them into a unified simulated dataset. The real dataset we chose is the IBD dataset (Gonzalez et al, 2022) studied the relationship between the gut microbiome and two main subtypes of inflammatory bowel disease (IBD): Crohn’s disease (CD) and ulcerative colitis (UC). It includes 108 CD and 66 UC samples, with profiles containing *p* = 5287 taxa.

We simulated two distinct datasets from the IBD dataset. We estimated separate location parameters for each of the two labels from the IBD dataset, denoted as *µ*_+_ ∈ R^5287^ and *µ*_−_ ∈ R^5287^. After that, we varied the location parameters to represent different levels of difficulty in distinguishing between labels, as follows:

- Setting 1: (2 ∗ *µ*_+_) and (2 ∗ *µ*_−_).
- Setting 2: Randomly select 10% of taxa with non-zero abundance and increase their values by 10%.

For each of these two simulated datasets, we generated the same number of positive and negative samples, matching the distribution of the IBD dataset.

Lastly, given that both the two simulated and seven real datasets have small sample sizes and high feature dimensionality, we included the third simulated dataset with a small sample size and low feature dimensionality to evaluate MIOSTONE’s performance in an atypical setting. Specifically, we employed the simulated dataset generated by Zhai et al (2024). The dataset comprises 48 genus aggregated from 2,964. We subsampled 50 samples, comprising 32 positive and 18 negative samples, respectively.

## A.1 Benchmark details

We evaluate the performance of MIOSTONE in comparison to nine baseline methods, divided into two categories: tree-agnostic methods and tree-aware methods. The former category comprises random for- est (RF), support vector machine (SVM) with a linear kernel, XGBoost (Chen and Guestrin, 2016), and multi-layer perceptron (MLP), while the latter includes, DeepBiome (Zhai et al, 2024), Ph-CNN (Fiora- vanti et al, 2018), PopPhy-CNN (Reiman et al, 2020), TaxoNN (Sharma et al, 2020), and MDeep (Wang et al, 2021). DeepBiome (Zhai et al, 2024) constructs its neural network architecture based on the phy- logenetic structure, where each hidden layer corresponds to a specific phylogenetic level (e.g., family, order, class, and phylum). The model is trained with a phylogenetic regularization technique using weight decay to prevent overfitting and improve generalization. Ph-CNN (Fioravanti et al, 2018) is based on a novel Phylo-Conv layer, which combines a convolutional operation with a neighbors detection algorithm. The network consists of a stack of Phylo-Conv layers, which are then flattened and followed by a fully connected (dense) layer, culminating in a final classification layer. PopPhy-CNN (Reiman et al, 2020) is a novel convolutional neural network designed to leverage the phylogenetic structure of microbial taxa for host phenotype prediction. The network processes a 2D matrix input, where the rows represent the phylogenetic tree and the columns contain the relative abundance of microbial taxa in a metagenomic sample. TaxoNN (Sharma et al, 2020) stratifies input taxa into different clusters based on their phylum information. The network comprises an ensemble of convolutional neural networks, each operating on a stratified cluster of taxa that share the same phylum. This approach is based on the rationale that taxa within the same phylum exhibit similarities and potential correlations, which can enhance predic- tive performance. MDeep (Wang et al, 2021) designs convolutional layers to mimic taxonomic ranks with multiple convolutional filters on each convolutional layer to capture the phylogenetic correlation among microbial species in a local receptive field and maintain the correlation structure across different convolutional layers via feature mapping.

We used the Scikit-learn implementation (Pedregosa et al, 2011) with default settings for the RF and SVM models. Specifically, the random forest classifier is trained with 100 trees, without a maximum tree depth constraint, and a minimum of 2 samples required to split an internal node. The support vector classifier is trained with a linear kernel, using L2 regularization with a coefficient of 1.0. The XGBoost classifier employs gradient boosting with decision trees, using a learning rate of 0.3, a maximum depth of 6, and 100 boosting rounds. For the tree-aware models, we used the recommended implementation settings. The MLP model is configured with a pyramid-shaped architecture with one hidden layer of size half of the input dimensionality. The DeepBiome model incorporates phylogenetic tree-based weight decay, utilizing Xavier uniform initialization without batch normalization or dropout. The Ph-CNN model consists of two 1D-CNN layers with 4 neighbors and 16 filters each, followed by a fully connected layer with 64 units and a dropout rate of 0.25. The PopPhy-CNN model features a 2D-CNN layer with 32 filters (kernel size: 3×10), followed by a fully connected layer with 512 units and a dropout rate of 0.3. We followed its built-in preprocessing pipeline, which involved log-scaling the normalized relative abundance of each taxon followed by subsequent min-max normalization. The TaxoNN model includes a 1D-CNN layer with 32 filters, another with 64 filters (both with a kernel size of 5), and a fully connected layer with 100 units. We utilized the recommended settings, *i.e.,* clustering the taxa based on their phylum information and ordering them according to Spearman correlation, as reported to yield optimal performance. The MDeep model comprises two 1D-CNN layers with 64 filters each and one layer with 32 filters, all with a kernel size of 8, a stride of 4, and a dropout rate of 0.5. All tree-aware models and MLP are trained for 200 epochs with a batch size of 512 to ensure convergence. During training, we used the AdamW optimizer with a learning rate of 0.001 and applied a cosine annealing scheduler. For a fair comparison, all models were trained and tested in the same environment: AMD EPYC 7302 16-Core Processor, NVIDIA RTX A6000 GPU, with 32GB DDR4 RAM.

## A.2 Detailed experiments

### A.2.1 Supervied learning

**Fig. A2.**
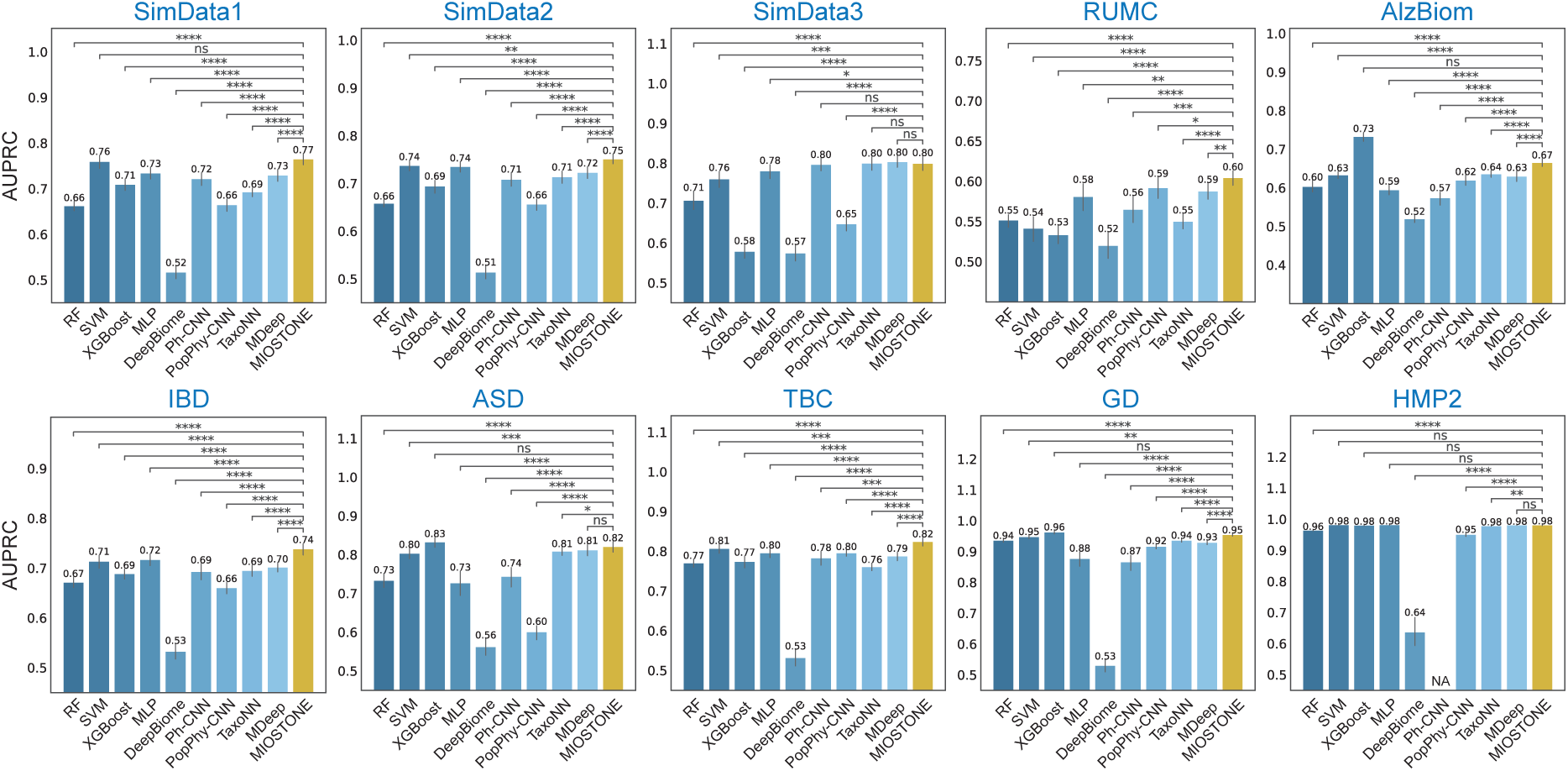
Performance of MIOSTONE in host’s disease status prediction in terms of AUROC. The evaluation was performed on three simulated and seven real microbiome datasets. MIOSTONE is compared against nine baseline methods, divided into two categories: tree-agnostic methods and tree-aware methods. Each model was trained by times using different train-test splits, and reported by the average performance along with 95% confidence intervals. The models’ performances are measured by the Area Under the Receiver Operating Characteristic Curve (AUROC) For scientific rigor, the performance comparison between MIOSTONE and any other baseline method is quantified using one-tailed two-sample t-tests to calculate p-values: ∗ ∗ ∗ ∗ p-value ≤ 0.0001; ∗ ∗ ∗p-value ≤ 0.001; ∗ ∗ p-value ≤ 0.01; ∗ : p-value ≤ 0.05; ns : p-value *>* 0.05.

### A.2.2 Transfer learning

**Fig. A3.**
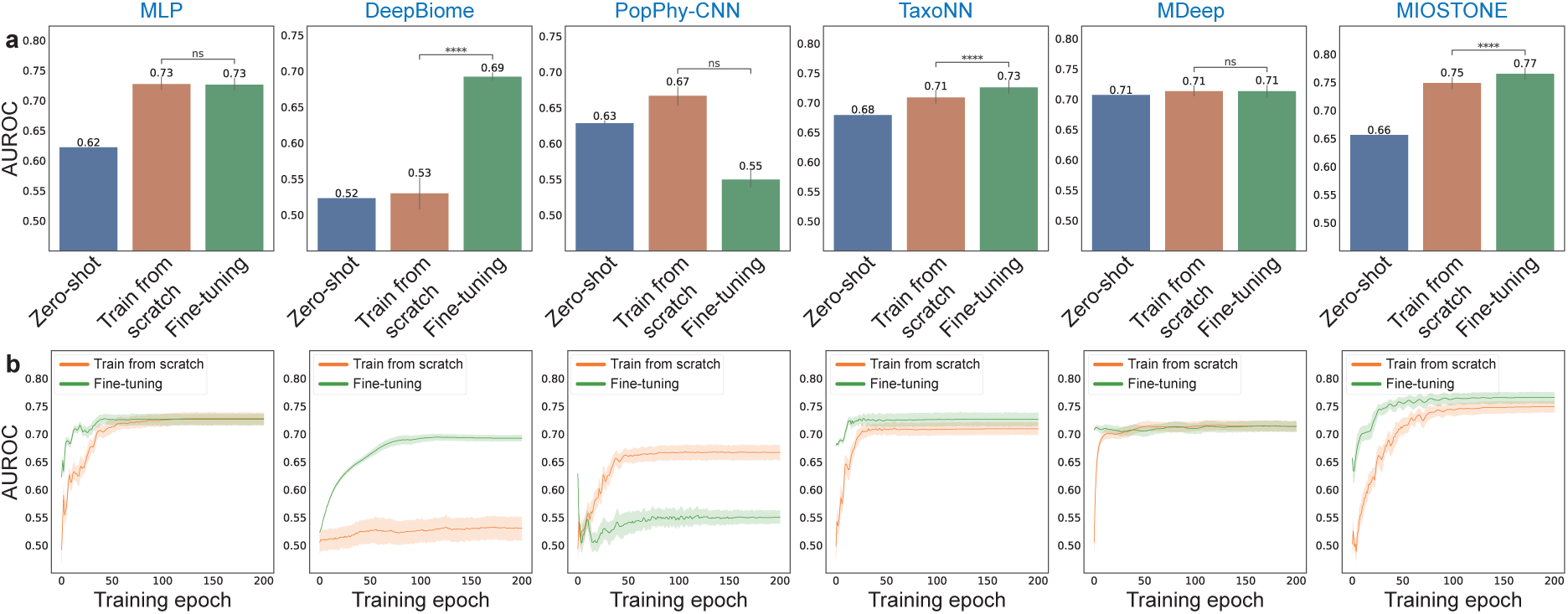
Performance of MIOSTONE in transferring knowledge from pre-trained models in terms of AUROC. (**a**) A model on the large HMP2 dataset is pre-trained and then employed for the smaller IBD dataset in three settings: direct prediction on IBD (*i.e.,* zero-shot), fine-tuning on IBD, and training IBD from scratch. Only tree-aware methods and MLP are included in the comparison, as most tree-agnostic methods are not well-suited for fine-tuning. Among the tree-aware methods, Ph-CNN is excluded because it is not scalable for processing the large HMP2 dataset. The prediction is conducted across three settings 20 times with varied train-test splits, and reported by the average performance assessed by AUROC, along with 95% confidence intervals. For scientific rigor, the performance between fine-tuning and training from scratch is quantified using one-tailed two-sample t-tests to calculate p-values. (**b**) The training dynamics of various models were evaluated by comparing fine-tuning with training from scratch, analyzing AUROC on test splits across different training epochs. MIOSTONE’s fine-tuning achieved better performance than training from scratch, requiring fewer training epochs.

### A.2.3 Representation learning

**Fig. A4.**
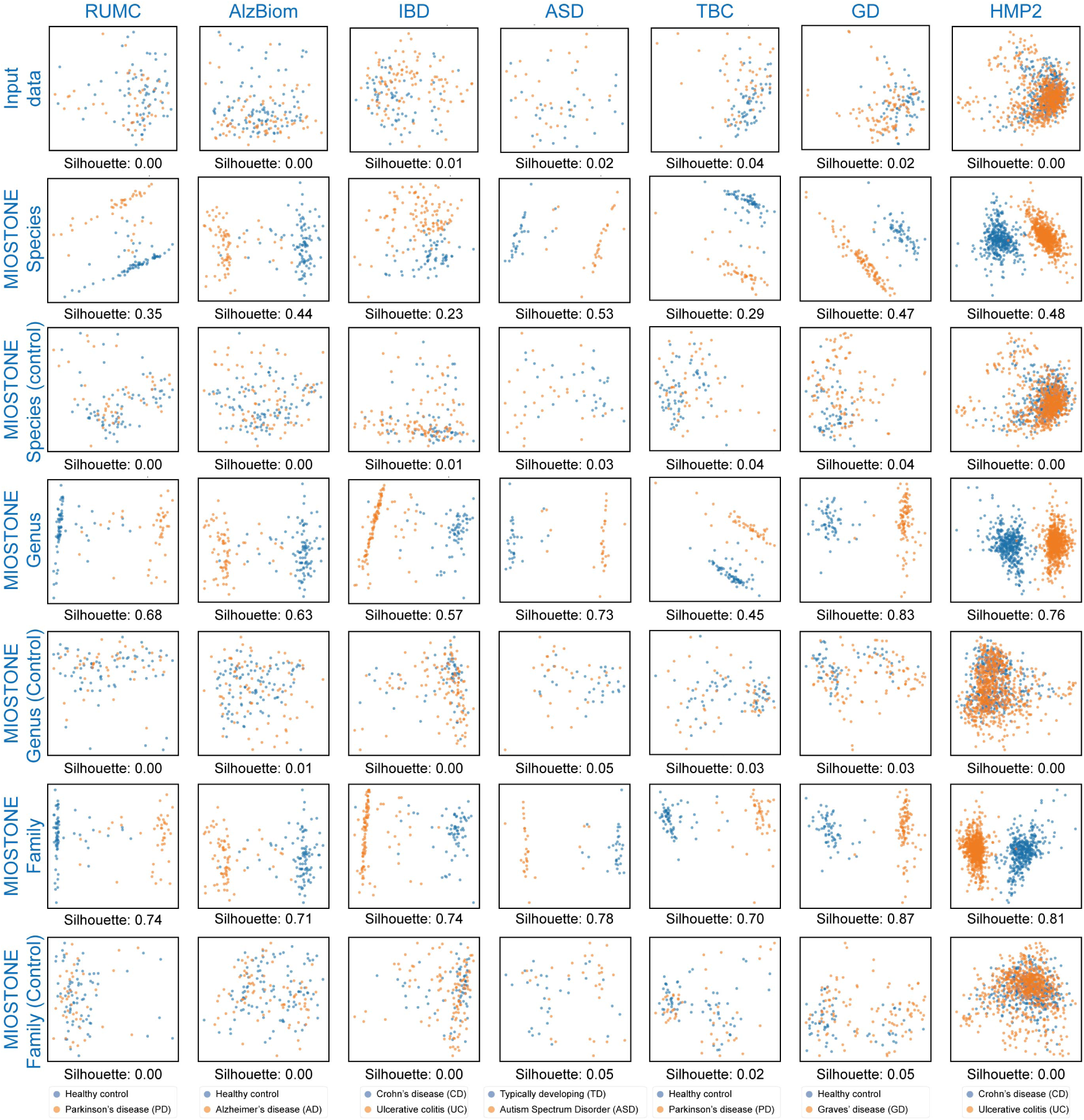
A sanity check of MIOSTONE’s internal neuron representations. We investigated whether MIOS- TONE’s internal neuron representations depend on the training data. If the representations were independent of the data and solely reliant on the model’s taxonomy-encoding architecture, it would be unreasonable and unreliable to draw convinc- ing conclusions from the model. As a sanity check, we used an untrained MIOSTONE model to project the internal neuron representations of samples onto a two-dimensional Principal Component space and evaluated their ability to distinguish between different disease subtypes, comparing the results to those of the trained MIOSTONE model. The untrained model exhibited no separation between disease subtypes, confirming that the model’s internal representations are data-dependent and capture disease-specific signatures during training.

### A.2.4 Model interpretation

**Fig. A5.**
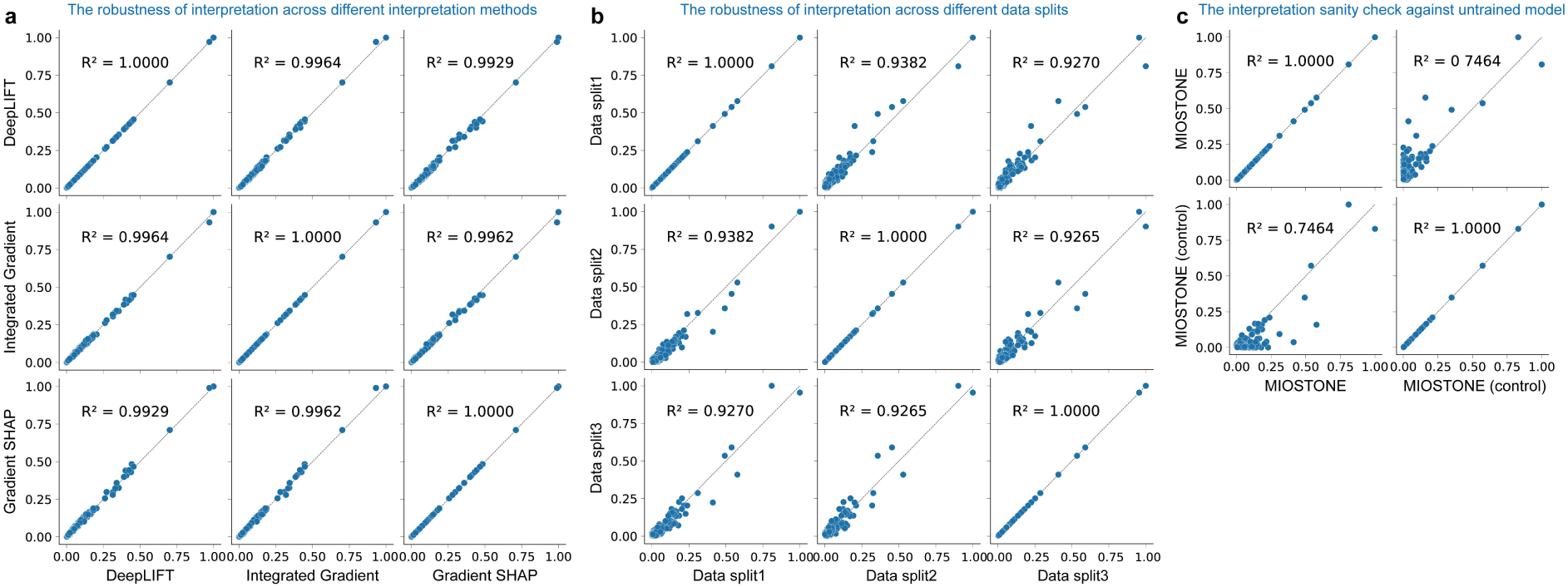
MIOSTONE is robust in discovering microbiome-disease associations using feature attribution methods. Feature attribution methods are used to interpret the MIOSTONE model and quantify the relationships between microbiome taxa and disease traits. (**a**) Three mainstream feature attribution methods–DeepLIFT, Integrated Gradients, and SHAP–demonstrate strong consistency in quantifying key microbiome-disease associations derived from the MIOS- TONE model. Therefore, the consistency of the methods makes it sufficient to present only the DeepLIFT results. (**b**) DeepLIFT demonstrates strong consistency in quantifying key microbiome-disease associations across different data splits. (**c**) A sanity check of MIOSTONE’s discovering microbiome-disease associations using feature attribution methods. We investigated whether the microbiome-disease associations reported by feature attribution methods depend on the training data. As a sanity check, we used an untrained MIOSTONE model and evaluated the consistency between the microbiome- disease associations it reported and those from a trained model. The low consistency suggests that the microbiome-disease associations reported by feature attribution methods are data-dependent.

### A.2.5 Control studies

**Fig. A6.**
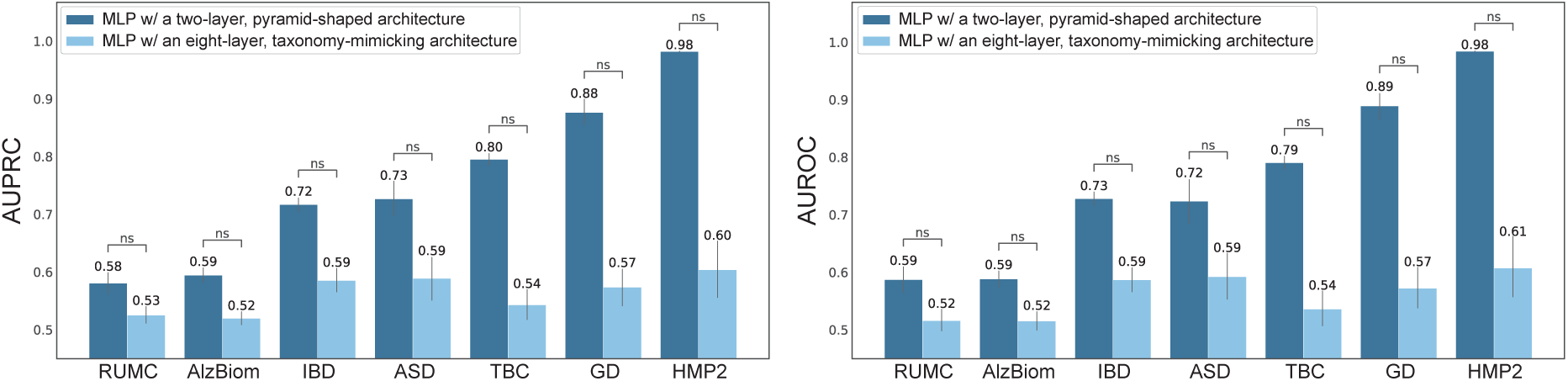
An MLP with a taxonomy-mimicking architecture does not enhance prediction accuracy. The MLP model used as baseline is configured with a pyramid-shaped architecture with one hidden layer of size half of the input dimensionality. Alternatively, we designed an MLP model that mirrors the taxonomy architecture, with each hidden layer corresponding to a specific taxonomic level, maintaining the same number of layers and neurons. The alternative MLP design significantly underperforms in prediction, as evidenced by lower AUPRC and AUROC scores. This suggests that taxonomy alone does not account for the superior predictive performance, as the increased number of parameters introduces additional challenges during training.

**Fig. A7.**
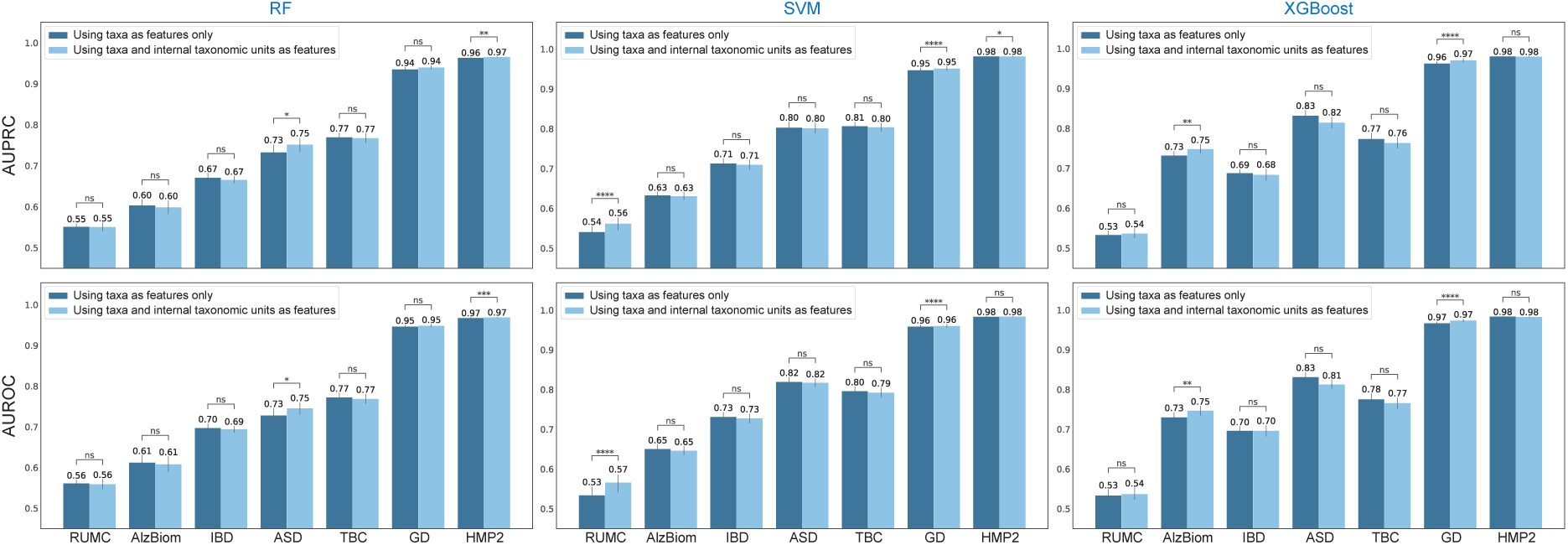
Flattening the internal taxonomic units as additional features does not effectively exploit the taxonomy. The feature value of each internal taxonomic unit is computed as the average of the values across all taxa within that specific taxonomic unit. Each tree-agnostic model takes as input the concatenated features from both the taxa and the internal taxonomic units. Treating the internal taxonomic units as additional features results in marginal or even worse predictive performance, as measured by both AUPRC and AUROC.

**Fig. A8.**
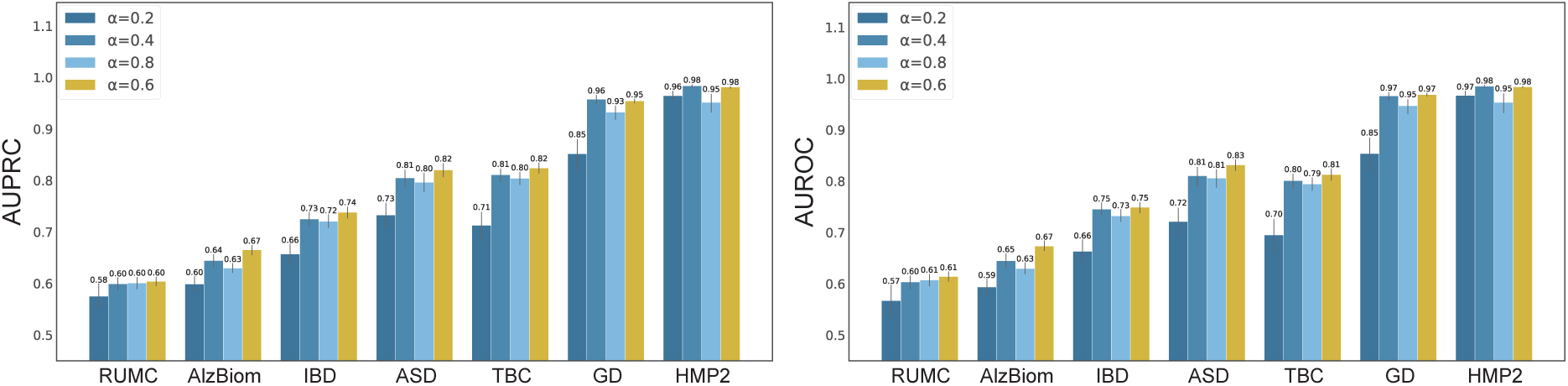
**MIOSTONE demonstrates robustness in selecting the hyperparameter that controls taxonomy- dependent representation dimensionality.**

**Fig. A9.**
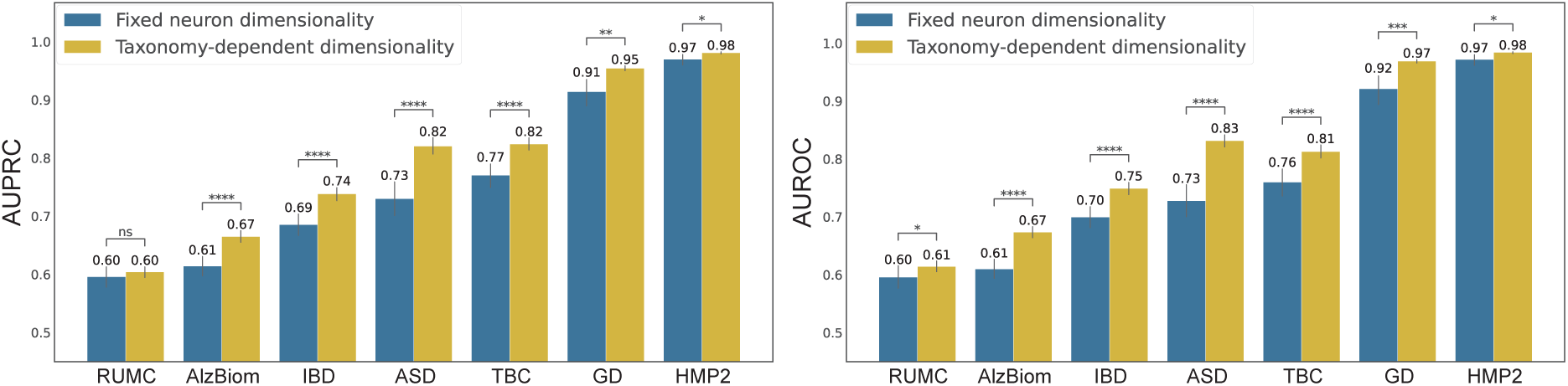
MIOSTONE demonstrates superior performance in using taxonomy-dependent representation dimensionality. MIOSTONE’s assigning larger taxonomic groups with greater representation dimensionality can aid in capturing more complex biological patterns to predict traits, compared to using fixed representation dimensionality.

**Fig. A10.**
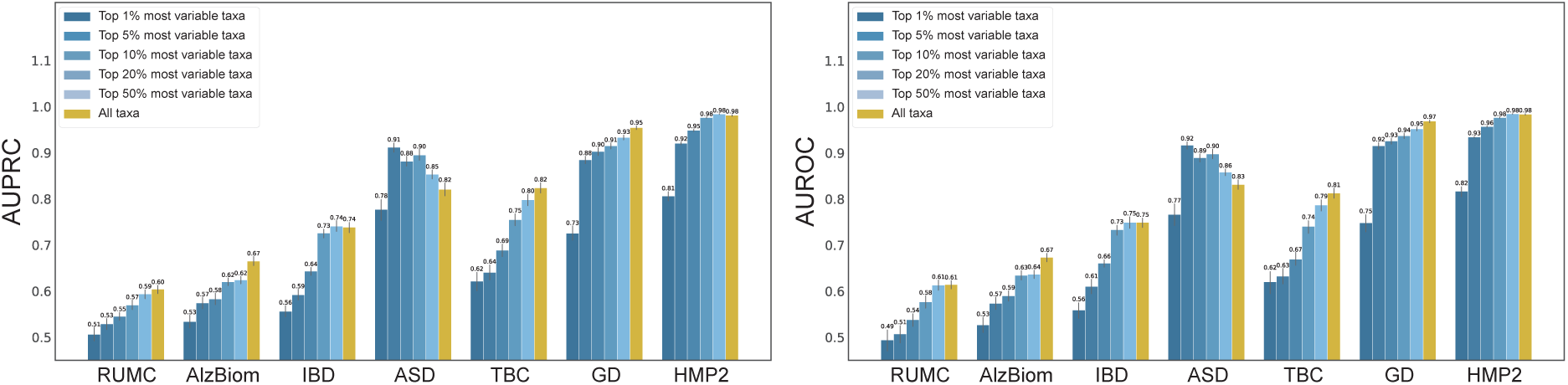
The curse of dimensionality cannot simply be mitigated using feature selection. MIOSTONE trained with all microbiome features, either outperforms or matches the performance of the model trained with a subset of highly variable taxa across most datasets.

**Fig. A11.**
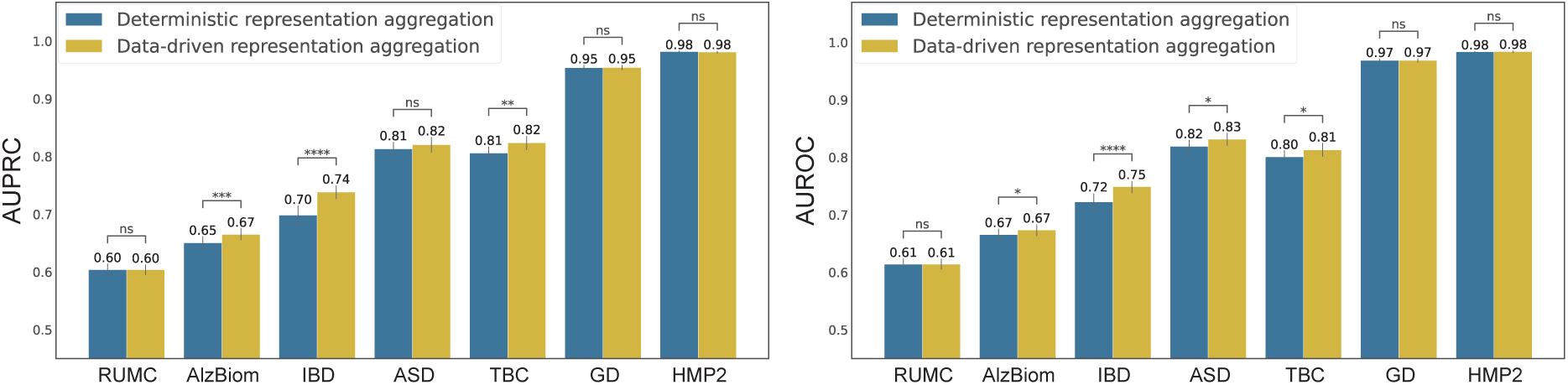
MIOSTONE demonstrates superior performance in using data-driven aggregation of neuron representations. MIOSTONE’s data-driven aggregation of neuron representations either outperforms or matches the performance of the deterministic selection of nonlinear representations across most datasets.

**Fig. A12.**
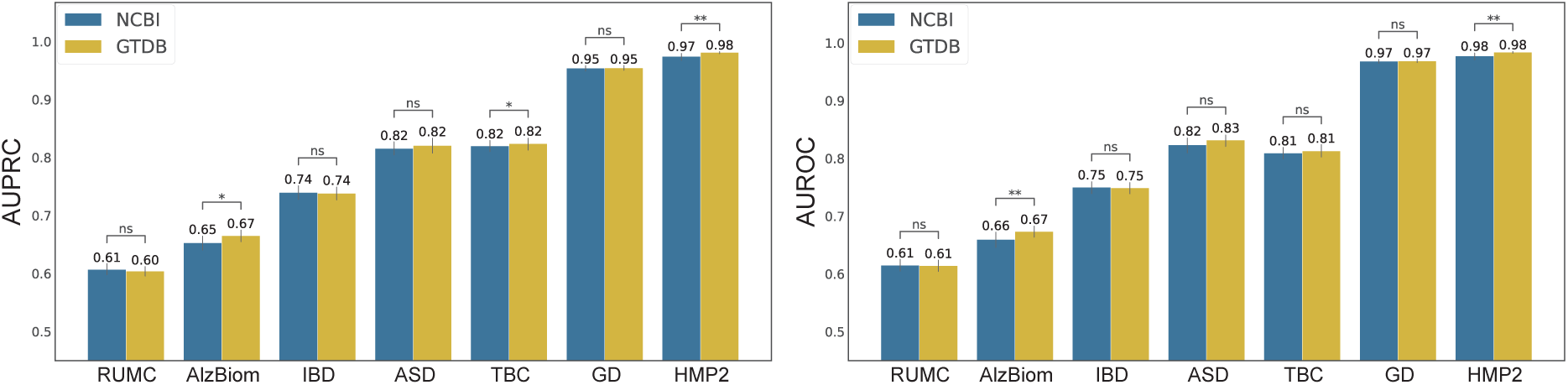
MIOSTONE demonstrates robustness across various taxonomic trees. Two variations of MIOSTONE utilizing taxonomies from GTDB and NCBI respectively, demonstrate comparable predictive performance across seven real microbiome datasets.

**Fig. A13.**
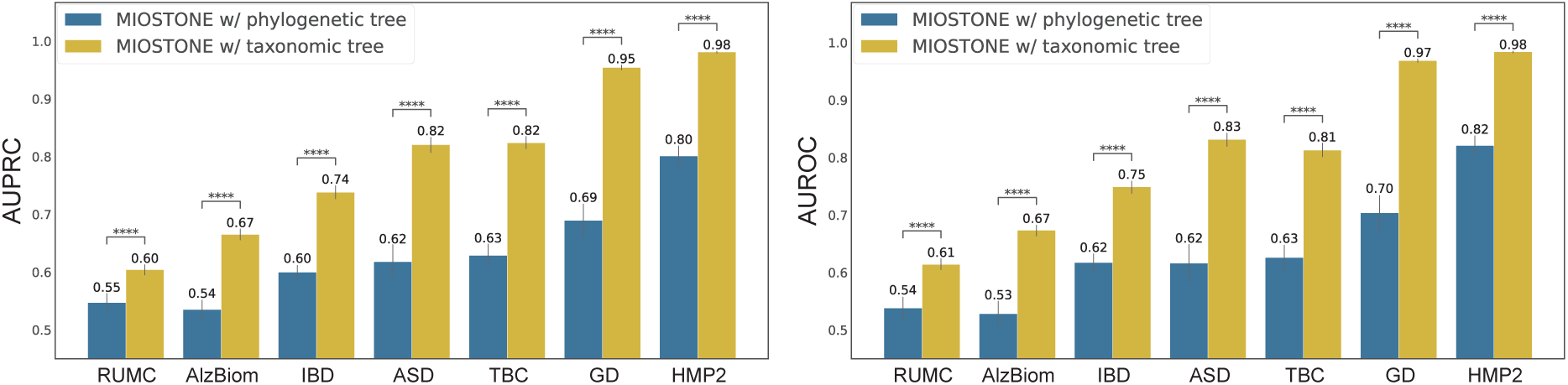
MIOSTONE demonstrates superior performance in using taxonomy-encoding architectures. While MIOSTONE can emulate any hierarchical correlation among taxa within its architecture, alternatives, such as phylogenic trees, perform significantly worse than the taxonomy-encoding architectures.

## Notes

### Competing Interest Statement

The authors have declared no competing interest.

### Summary of Updates

The paper has been updated with new experiment results, revised figures, and many other improvements.

## References

1. Adebayo J, Gilmer J, Muelly M, et al (2018) Sanity checks for saliency maps. Advances in Neural Information Processing Systems 31

2. Aitchison J (1982) The statistical analysis of compositional data. Journal of the Royal Statistical Society: Series B (Methodological) 44(2):139–160

3. Alam MT, Amos GC, Murphy AR, et al (2020) Microbial imbalance in inflammatory bowel disease patients at different taxonomic levels. Gut Pathogens 12:1–8

4. Albanese D, Filippo CD, Cavalieri D, et al (2015) Explaining diversity in metagenomic datasets by phylogenetic-based feature weighting. PLoS Computational Biology 11(3):e1004186

5. Andrews TS, Hemberg M (2018) False signals induced by single-cell imputation. F1000Research 7

6. Armstrong G, Martino C, Morris J, et al (2022) Swapping metagenomics preprocessing pipeline components offers speed and sensitivity increases. mSystems 7(2):e01378–21

7. Barber RF, Candèes EJ (2015) Controlling the false discovery rate via knockoffs. The Annals of Statistics 43(5):2055–2085

8. Bengio Y, Courville A, Vincent P (2013) Representation learning: A review and new perspectives. IEEE Transactions on Pattern Analysis and Machine Intelligence 35(8):1798–1828

9. Boktor J, Sharon G, Metman LV, et al (2023) Integrated multi-cohort analysis of the Parkinson’s disease gut metagenome. Movement Disorders 38(3):399–409

10. Chen T, Guestrin C (2016) XGBoost: A scalable tree boosting system. In: ACM SIGKDD International Conference on Knowledge Discovery and Data Mining. ACM, New York, NY, USA, KDD ’16, pp 785–794

11. Chen W, Jiang Y, Noble WS, et al (2024a) Error-controlled non-additive interaction discovery in machine learning models. arXiv preprint arXiv:240817016

12. Chen W, Noble WS, Lu YY (2024b) DeepROCK: Error-controlled interaction detection in deep neural networks. Neural Information Processing Systems (Intepretable AI Workshop)

13. Dan Z, Mao X, Liu Q, et al (2020) Altered gut microbial profile is associated with abnormal metabolism activity of Autism Spectrum Disorder. Gut Microbes 11(5):1246–1267

14. Ditzler G, Morrison JC, Lan Y, et al (2015) Fizzy: feature subset selection for metagenomics. BMC Bioinformatics 16:1–8

15. Elmarakeby HA, Hwang J, Arafeh R, et al (2021) Biologically informed deep neural network for prostate cancer discovery. Nature 598(7880):348–352

16. Finucane MM, Sharpton TJ, Laurent TJ, et al (2014) A taxonomic signature of obesity in the microbiome? Getting to the guts of the matter. PloS One 9(1):e84689

17. Fioravanti D, Giarratano Y, Maggio V, et al (2018) Phylogenetic convolutional neural networks in metagenomics. BMC Bioinformatics 19:1–13

18. Gonzalez A, Navas-Molina JA, Kosciolek T, et al (2018) Qiita: rapid, web-enabled microbiome meta- analysis. Nature Methods 15(10):796–798

19. Gonzalez CG, Mills RH, Zhu Q, et al (2022) Location-specific signatures of Crohn’s disease at a multi- omics scale. Microbiome 10(1):133

20. Grice EA, Segre JA (2012) The human microbiome: Our second genome. Annual Review of Genomics and Human Genetics 13:151–170

21. Han S, Pool J, Tran J, et al (2015) Learning both weights and connections for efficient neural network. In: Advances in Neural Information Processing Systems

22. Hao Y, Hao S, Andersen-Nissen E, et al (2021) Integrated analysis of multimodal single-cell data. Cell

23. He M, Zhao N, Satten GA (2024) MIDASim: a fast and simple simulator for realistic microbiome data. Microbiome 12(1):135

24. Ioffe S, Szegedy C (2015) Batch normalization: Accelerating deep network training by reducing internal covariate shift. International Conference on Machine Learning pp 448–456

25. Iuchi H, Matsutani T, Yamada K, et al (2021) Representation learning applications in biological sequence analysis. Computational and Structural Biotechnology Journal 19:3198–3208

26. Jiang R, Li WV, Li JJ (2021) mbImpute: an accurate and robust imputation method for microbiome data. Genome Biology 22(1):192

27. Jiang R, Sun T, Song D, et al (2022) Statistics or biology: the zero-inflation controversy about scRNA-seq data. Genome Biology 23(1):31

28. Kabeerdoss J, Jayakanthan P, Pugazhendhi S, et al (2015) Alterations of mucosal microbiota in the colon of patients with inflammatory bowel disease revealed by real time polymerase chain reaction amplification of 16s ribosomal ribonucleic acid. Indian Journal of Medical Research 142(1):23–32

29. Kharchenko PV (2021) The triumphs and limitations of computational methods for scRNA-seq. Nature Methods 18(7):723–732

30. Knights D, Parfrey LW, Zaneveld J, et al (2011) Human-associated microbial signatures: examining their predictive value. Cell Host & Microbe 10(4):292–296

31. Kojima A, Nakano K, Wada K, et al (2012) Infection of specific strains of streptococcus mutans, oral bacteria, confers a risk of ulcerative colitis. Scientific Reports 2(1):332

32. Laske C, Müller S, Preische O, et al (2022) Signature of Alzheimer’s disease in intestinal microbiome: Results from the AlzBiom study. Frontiers in Neuroscience 16:792996

33. Lee AA, Rao K, Limsrivilai J, et al (2020) Temporal gut microbial changes predict recurrent clostrid- iodes difficile infection in patients with and without ulcerative colitis. Inflammatory Bowel Diseases 26(11):1748–1758

34. Li B, Zhong D, Jiang X, et al (2021) TopoPhy-CNN: integrating topological information of phylogenetic tree for host phenotype prediction from metagenomic data. In: IEEE International Conference on Bioinformatics and Biomedicine (BIBM), IEEE, pp 456–461

35. Linderman GC, Zhao J, Roulis M, et al (2022) Zero-preserving imputation of single-cell RNA-seq data. Nature Communications 13(1):192

36. Liu B, Wei Y, Zhang Y, et al (2017) Deep neural networks for high dimension, low sample size data. In: International Joint Conference on Artificial Intelligence, pp 2287–2293

37. Lloyd-Price J, Arze C, Ananthakrishnan AN, et al (2019) Multi-omics of the gut microbial ecosystem in inflammatory bowel diseases. Nature 569(7758):655–662

38. Loshchilov I, Hutter F (2019) Decoupled weight decay regularization. International Conference on Learning Representations

39. Louizos C, Welling M, Kingma DP (2018) Learning sparse neural networks through l 0 regularization. International Conference on Learning Representations

40. Lu YY, Fan Y, Lv J, et al (2018) DeepPINK: reproducible feature selection in deep neural networks. In: Advances in Neural Information Processing Systems

41. Lundberg SM, Lee SI (2017) A unified approach to interpreting model predictions. Advances in Neural Information Processing Systems

42. Ma J, Yu MK, Fong S, et al (2018) Using deep learning to model the hierarchical structure and function of a cell. Nature Methods 15(4):290–298

43. Medina RH, Kutuzova S, Nielsen KN, et al (2022) Machine learning and deep learning applications in microbiome research. ISME Communications 2(1):98

44. Mills RH, Dulai PS, Váazquez-Baeza Y, et al (2022) Multi-omics analyses of the ulcerative colitis gut microbiome link bacteroides vulgatus proteases with disease severity. Nature Microbiology 7(2):262–276

45. Parks DH, Chuvochina M, Waite DW, et al (2018) A standardized bacterial taxonomy based on genome phylogeny substantially revises the tree of life. Nature Biotechnology 36(10):996–1004

46. Pasolli E, Truong DT, Malik F, et al (2016) Machine learning meta-analysis of large metagenomic datasets: tools and biological insights. PLoS Computational Biology 12(7):e1004977

47. Paszke A, Gross S, Massa F, et al (2019) PyTorch: An imperative style, high-performance deep learning library. In: Advances in Neural Information Processing Systems. Curran Associates, Inc., Vancouver, Canada, p 8024–8035

48. Pedregosa F, Varoquaux G, Gramfort A, et al (2011) Scikit-learn: Machine learning in Python. Journal of Machine Learning Research 12:2825–2830

49. Qin J, Li R, Raes J, et al (2010) A human gut microbial gene catalogue established by metagenomic sequencing. Nature 464(7285):59–65

50. Qin J, Li Y, Cai Z, et al (2012) A metagenome-wide association study of gut microbiota in type 2 diabetes. Nature 490(7418):55–60

51. Reiman D, Metwally AA, Sun J, et al (2020) PopPhy-CNN: a phylogenetic tree embedded architecture for convolutional neural networks to predict host phenotype from metagenomic data. IEEE Journal of Biomedical and Health Informatics 24(10):2993–3001

52. Rousseeuw PJ (1987) Silhouettes: a graphical aid to the interpretation and validation of cluster analysis. Journal of Computational and Applied Mathematics 20:53–65

53. Sasaki K, Inoue J, Sasaki D, et al (2019) Construction of a model culture system of human colonic microbiota to detect decreased Lachnospiraceae abundance and butyrogenesis in the feces of ulcerative colitis patients. Biotechnology Journal 14(5):1800555

54. Sekirov I, Russell SL, Antunes CM, et al (2010) Gut microbiota in health and disease. Physiological Reviews

55. Sharma D, Paterson AD, Xu W (2020) TaxoNN: ensemble of neural networks on stratified microbiome data for disease prediction. Bioinformatics 36(17):4544–4550

56. Shrikumar A, Greenside P, Shcherbina A, et al (2017) Learning important features through propagating activation differences. In: International Conference on Machine Learning

57. Shtossel O, Isakov H, Turjeman S, et al (2023) Ordering taxa in image convolution networks improves microbiome-based machine learning accuracy. Gut Microbes 15(1):2224474

58. Storey JD (2003) The positive false discovery rate: A bayesian interpretation and the q-value. The Annals of Statistics 31(6):2013–2035

59. Stuart T, Butler A, Hoffman P, et al (2019) Comprehensive integration of single-cell data. Cell 77(7):1888–1902

60. Sundararajan M, Taly A, Yan Q (2017) Axiomatic attribution for deep networks. In: International Conference on Machine Learning

61. Tsang M, Cheng D, Liu Y (2018) Detecting statistical interactions from neural network weights. International Conference on Learning Representations

62. Turnbaugh PJ, Ley RE, Mahowald MA, et al (2006) An obesity-associated gut microbiome with increased capacity for energy harvest. Nature 444(7122):1027–1031

63. Vogt NM, Kerby RL, Dill-McFarland KA, et al (2017) Gut microbiome alterations in Alzheimer’s disease. Scientific Reports 7(1):13537

64. Wang Y, Bhattacharya T, Jiang Y, et al (2021) A novel deep learning method for predictive modeling of microbiome data. Briefings in Bioinformatics 22(3):bbaa073

65. Washburne AD, Morton JT, Sanders J, et al (2018) Methods for phylogenetic analysis of microbiome data. Nature Microbiology 3(6):652–661

66. Weiss K, Khoshgoftaar TM, Wang D (2016) A survey of transfer learning. Journal of Big data 3(1):1–40

67. Wolf FA, Angerer P, Theis FJ (2018) SCANPY: large-scale single-cell gene expression data analysis. Genome Biology 19:1–5

68. Xiao J, Chen L, Johnson S, et al (2018) Predictive modeling of microbiome data using a phylogeny- regularized generalized linear mixed model. Frontiers in Microbiology 9:1391

69. Ye SH, Siddle KJ, Park DJ, et al (2019) Benchmarking metagenomics tools for taxonomic classification. Cell 178(4):779–794

70. Zeng Y, Li J, Wei C, et al (2022) mbDenoise: microbiome data denoising using zero-inflated probabilistic principal components analysis. Genome Biology 23(1):94

71. Zhai J, Choi Y, Yang X, et al (2024) DeepBiome: a phylogenetic tree informed deep neural network for microbiome data analysis. Statistics in Biosciences pp 1–25

72. Zhu Q, Mai U, Pfeiffer W, et al (2019) Phylogenomics of 10,575 genomes reveals evolutionary proximity between domains Bacteria and Archaea. Nature Communications 10(1):5477

73. Zhu Q, Hou Q, Huang S, et al (2021) Compositional and genetic alterations in Graves’ disease gut microbiome reveal specific diagnostic biomarkers. The ISME journal 15(11):3399–3411

74. Zhu Q, Huang S, Gonzalez A, et al (2022) Phylogeny-aware analysis of metagenome community ecology based on matched reference genomes while bypassing taxonomy. mSystems 7(2):e00167–22

